# Topologically Engineered Tetrahedral Antibody Architectures for Multispecific Therapy

**DOI:** 10.1101/2022.03.23.485397

**Authors:** Daniel J. Capon, Larisa A. Troitskaya, Marina E. Fomin, Malcolm L. Gefter, Ursula Edman, Brendon Frank, Benjamin Z. Capon, Jing Jin, Rachel Martinelli, Ginger A. Ferguson, Ramona Vestweber, Cheila Rocha, Berit Roshani, Victor Corman, Christian Drosten, Nadine Krüger, Stefan Pöhlmann, Graham Simmons, Nelson L.S. Chan

## Abstract

Traditional Y-shaped IgG topologies restrict conventional multispecific antibodies by imposing steric constraints that hinder target binding and stability. To overcome this, we engineered tetrahedral antibodies, a multispecific class that non-covalently self-assembles via two hinge-integrated collectrin-like domains (CLDs). This non-planar, tetrapartite architecture coordinates up to six unobstructed Fab, Fc, and extracellular domains. Parallel to the clinical translation of our dual-Fc anti-CD19/CD20 tetrahedral antibody for systemic lupus erythematosus (SLE), we evaluated an anti-SARS-CoV-2 candidate that exhibited in vivo efficacy in non-human primates. Furthermore, we demonstrate functional FcγR binding cooperativity across the dual-Fc domains of anti-CD19/CD20 tetrahedral antibodies, while introducing an interface disulfide bond into the CLD dimer increases structural stability as evidenced by effectively abolishing CLD dynamic dissociation and resolving the dissociation equilibrium into discrete, stable quaternary states. These findings establish tetrahedral architectures as a robust, translatable multispecific platform.

---

The native Y-shaped antibody represents an elegant structural archetype. Within this dimeric geometry, two heavy and two light chains are assembled via a disulfide-bonded hinge region to ensure thermodynamic stability and unobstructed ligand engagement by the Fab and Fc domains. By contrast, most engineered multispecific immunoglobulin formats utilize classical linear concatenation strategies, fusing accessory binding domains directly to the amino or carboxy termini of native antibody chains.^1,2^ These complex, multi-domain architectures frequently suffer from compromised expression yields, mispairing, and severe steric hindrance, which collectively restrict accessibility to both target antigens and immune effector receptors.^3,4^

Beyond steric crowding at individual chain termini, the primary biophysical constraint of conventional multispecific antibodies is their strict planar geometry. The rigid, two-dimensional layout of standard immunoglobulin scaffolds limits the spatial orientation, rotational freedom, and flexibility of the binding arms. When attempting to bridge distinct cells, such as immunological synapses between effector cells and target pathogens, or cooperatively bind dense, clustered cell-surface antigens, a planar configuration restricts optimal alignment of all binding domains. This geometric limitation introduces a significant thermodynamic penalty, resulting in compromised binding cooperativity, incomplete receptor occupancy, and sub-optimal therapeutic efficacy.

Nature frequently overcomes such geometric constraints through complex quaternary structures that self-assemble via non-covalent forces to achieve precise three-dimensional spatial orientations.^5,6^ Inspired by these evolutionary mechanisms, we moved beyond traditional linear fusion strategies by leveraging non-covalent cross-dimerization to engineer a three-dimensional antibody topology.^7,8^

Here, we describe a multispecific platform utilizing the collectrin-like domain (CLD) derived from human angiotensin-converting enzyme 2 (ACE2),^9,10^ integrated directly into the antibody hinge region. Acting in concert with the adjacent Fc domain, these integrated modules serve as a structural nucleus, directing the self-assembly of two to four distinct polypeptide chains into a highly stable, symmetric tetrahedral architecture. This non-planar layout projects its binding faces outward into three-dimensional space, decoupling multivalent domain assembly from steric interference. By ensuring unobstructed spatial access for up to six individual Fab, Fc, or extracellular domains, this tetrahedral architecture enables enhanced, cooperative binding across complex cellular and antigenic landscapes.

## Results

### Discovery and Structural Characterization of Tetrahedral Assemblies

To systematically evaluate how distinct domain interactions dictate molecular topology, we engineered a series of constructs governed by varying dimerization forces (**Figs. 1a-d**). Structural analysis revealed four topologically distinct types of ACE2-Fc dimers: those driven solely by the Fc domain (**Fig. 1a**), co-driven cooperatively by both the Fc domain and CLD dimer (**Fig. 1b-c**), and solely by the CLD dimer (**Fig. 1d**).

**Fig. 1.**
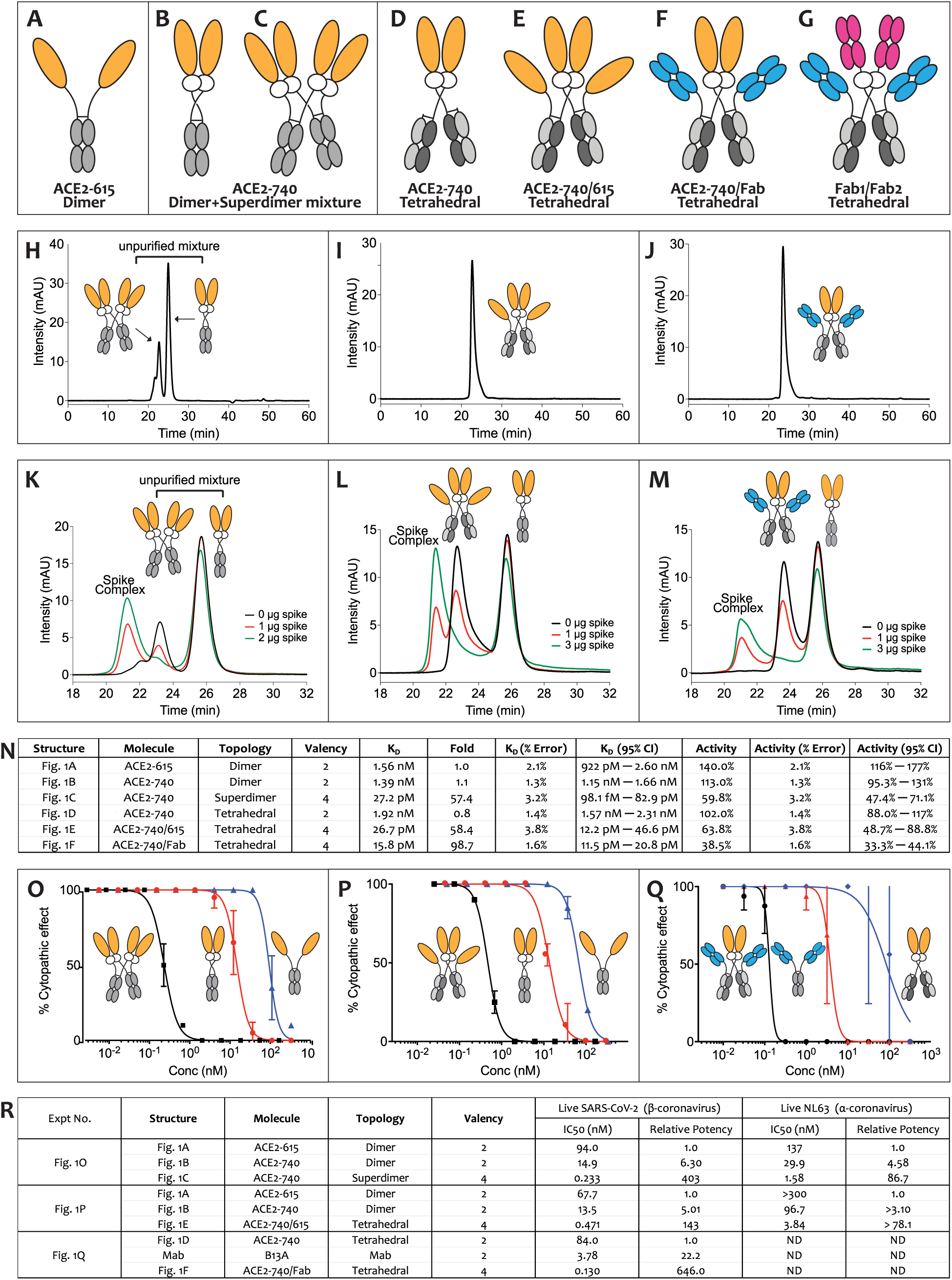
**Engineering tetrahedral architectures via CLD-driven dimerization provides structural homogeneity, cooperative SARS-CoV-2 spike binding, and orders of magnitude enhanced live virus neutralization**. (a–g) Schematics illustrating (a) the baseline Fc-driven ACE2-615 dimer, (b, c) the co-secreted ACE2-740 dimer/superdimer mixture driven by cross-dimerization of the Fc domain and the CLD dimer, and (d–g) tetrahedral antibodies driven solely by single CLD dimer, specifically (d) the ACE2-740 core tetrahedral antibody with four domains, and (e-g) elaborated tetravalent versions of the core architecture with six domains, specifically, (e) the tetravalent monospecific ACE2-740/615 tetrahedral antibody, (f) the tetravalent bispecific ACE2-740/Fab tetrahedral antibody, and (g) the tetravalent bispecific Fab1/Fab2 tetrahedral antibody. White circles represent the CLD core; orange ovals indicate the ACE2 peptidase; blue and magenta ovals denote distinct Fab domains; and gray structures indicate Fc heterodimers. (h– j) SE-HPLC profiles demonstrating (h) ACE2-740 dimer and superdimer are co-secreted as a topological mixture, whereas the optimized (i) ACE2-740/615 and (j) ACE2-740/B13A tetrahedral antibodies are expressed and secreted as a single, homogenous species. (k–m) Stoichiometric competition binding demonstrating cooperative spike binding favoring ACE2-740 superdimer and the ACE2-740/615 and ACE2-740/Fab tetrahedral antibodies over ACE2-740 dimer, monitoring complex formation with individual SARS-CoV-2 spike protein trimers across a titration range of 0 to 2 μg using (k) the unpurified mixture of ACE2-740 dimer and superdimer, (l) ACE2-740/615 tetrahedral antibody mixed with ACE2-740 dimer, and (m) ACE2-740/Fab tetrahedral antibody mixed with ACE2-740 dimer. (n) Solution-binding summary table detailing molecule configurations, topological assignments, calculated valence, equilibrium dissociation constants (K_D_), error percentages, and absolute activity values derived via KinExA modeling. (o–q) Live virus neutralization dose-response curves plotting percentage cytopathic effect (CPE) suppression of (o) ACE2-740 superdimer, and (p) ACE2-740/615 tetrahedral antibody, and (q) ACE2-740/Fab tetrahedral antibody. (r) Summary table detailing half-maximal inhibitory concentrations (*IC₅₀*) and relative neutralization potencies for live SARS-CoV-2 and human coronavirus NL63 (shown in Supplementary **Figs. 5a**, b).

While the ACE2-615 dimer (**Fig. 1a**) and ACE2-740 tetrahedral construct (**Fig. 1d**) were each resolved as a single, homogenous species, the third construct, co-driven by both the Fc and CLD modules (**Fig.1** **b, c**), existed as a mixture of two topologically distinct forms (**Fig. 1h**) as shown by size-exclusion high-performance liquid chromatography (SE-HPLC). We confirmed these distinct structural conformations using site-specific enzymatic cleavage by IdeZ protease followed by multi-angle light scattering coupled with size-exclusion chromatography (SEC-MALS) (**Supplementary Figs. 1–2**, **Supplementary Table 1**).

SEC-MALS analysis of this mixture (**Supplementary Fig. 3**) indicated that the smaller species corresponded to the predicted closed ACE2-740 dimer (**Fig. 1b**), whereas the larger species represented a dimer-of-dimers, designated as ACE2-740 “superdimer” (**Fig. 1c**). This novel quaternary architecture arises from the reciprocal cross-dimerization of two pairs of CLD-dimerizing polypeptides and two pairs of Fc-dimerizing polypeptides. Structural modeling suggests this superdimeric antibody adopts a rigid tetrahedral-like configuration, projecting its two ACE2 peptidase dimers and two Fc dimers outward into three-dimensional space from a centralized CLD nucleus.

### Cooperative Binding and Structural Optimization of Tetrahedral Antibodies

To evaluate the functional consequences of the unique three-dimensional topology of the tetrahedral antibody, we conducted stoichiometric competition binding studies. When unpurified mixtures of the ACE2-740 dimer (**Fig. 1b**) and ACE2-740 superdimer (**Fig. 1c**) were titrated with limiting amounts of aggregate-free SARS-CoV-2 spike protein trimers, we observed a hierarchical order of binding with superdimer–spike complexes formed preferentially over dimer–spike complexes, indicating a pronounced thermodynamic preference for superdimer binding (**Fig. 1k**). Identical competitive advantages were observed when testing equimolar mixtures of the purified ACE2-740 superdimer against all three ACE2 dimer configurations (**Supplementary Fig. 4**).

While the dimeric cross-dimerization platform yielded a mixture of topologies, we achieved structural homogeneity through targeted molecular engineering. By restricting the constructs to a single pair of CLD-dimerizing polypeptides, we guided protein assembly into a single, defined topological form (**Fig. 1i-j**). These optimized, symmetrically paired architectures are collectively designated as tetrahedral antibodies (**Fig. 1d-g**). This modular platform accommodates various combinations of variable regions or Fc-fusion domains. Notably, the bispecific tetrahedral configurations (**Figs. 1f-g**) can be engineered in two alternative spatial orientations depending on which binding moiety is directly fused to the CLD nucleus versus the Fc domain—a structural variation that can exert significant functional effects.

Using this design strategy, we expressed the tetravalent monospecific ACE2-740/615 tetrahedral antibody (**Fig. 1e**) and the tetravalent bispecific ACE2-740/B13A tetrahedral antibody (**Fig. 1f**). SE-HPLC confirmed that both tetrahedral antibodies were secreted strictly as single, homogenous molecular species (**Figs. 1i-j**). In stoichiometric titrations, both the ACE2-740/615 and ACE2-740/B13A tetrahedral antibodies demonstrated high-affinity binding to individual spike trimers, comparable to the ACE2-740 superdimer (**Figs. 1l-m**). Quantitative solution-binding analyses by KinExA (**Fig. 1n**) validated these observations. The standard planar constructs, ACE2-615 dimer and ACE2-740 dimer, as well as the bivalent ACE2-740 tetrahedral antibody, bound the spike protein with equilibrium dissociation constants (K_D_) of 1.56 nM, 1.39 nM, and 1.92 nM, respectively. By contrast, all three tetravalent tetrahedral architectures, the ACE2-740 superdimer, and the ACE2-740/615 and ACE2-740/B13A tetrahedral antibodies, exhibited K_D_ values of 27.2 pM, 26.7 pM, and 15.8 pM, respectively. These metrics establish a 57- to 99-fold cooperative binding advantage for the tetravalent tetrahedral antibodies over traditional planar counterparts.

### Potent Viral Neutralization and Spike Trimer Binding

To evaluate the therapeutic potential of these tetrahedral architectures against pathogens utilizing the ACE2 entry receptor, we benchmarked their neutralization capacity against live SARS-CoV-2 (**Figs. 1o-q**) and live human beta-coronavirus NL63 (HCoV-NL63) (**Supplementary Fig. 5**). The neutralization potencies (**Fig. 1r**) directly mirrored the trends observed in our solution-binding kinetics (**Fig. 1n**). Against SARS-CoV-2, the ACE2-740 superdimer and ACE2-740/615 tetrahedral antibody demonstrated 403-fold and 143-fold greater potency than the standard planar ACE2-615 dimer, respectively; against HCoV-NL63, they exhibited 87-fold and greater than 78-fold potency enhancements. Similarly, the bispecific ACE2-740/B13A tetrahedral antibody displayed a 646-fold and 29-fold increase in neutralization potency against SARS-CoV-2 compared to the ACE2-740 dimer and the parental B13A monoclonal antibody,^11^ respectively.

This neutralization advantage was corroborated by cell-surface receptor-binding inhibition assays (**Supplementary Fig. 5**, **Supplementary Table 2**). The ACE2-740 superdimer and ACE2-740/615 tetrahedral antibody blocked spike trimer binding to cell-surface ACE2 with an 8.9-fold and 10.6-fold cooperative advantage over the ACE2-740 dimer, respectively. Furthermore, the ACE2- 740/B13A tetrahedral antibody blocked receptor binding with an 11.2-fold and 3.8-fold advantage over the ACE2-740 dimer and parental B13A antibody, respectively.

### Enhancing Monoclonal Antibody Potency via Tetrahedral Formatting

To demonstrate the generalizability of the tetrahedral architecture for enhancing the potency of conventional therapeutic immunoglobulins, we engineered a panel of tetravalent monospecific Fab-Fab tetrahedral antibodies (**Fig. 2a**) that incorporated the Fab domains from eight clinically FDA-authorized or approved antibodies.^12–16^ Stoichiometric competition binding experiments with individual spike trimers confirmed that these reformatted tetrahedral antibodies formed stable complexes with target spikes in high thermodynamic preference over their bivalent monospecific parental antibody counterparts (**Figs. 2c-j**, **Supplementary Figs. 6, 7**).

**Fig. 2.**
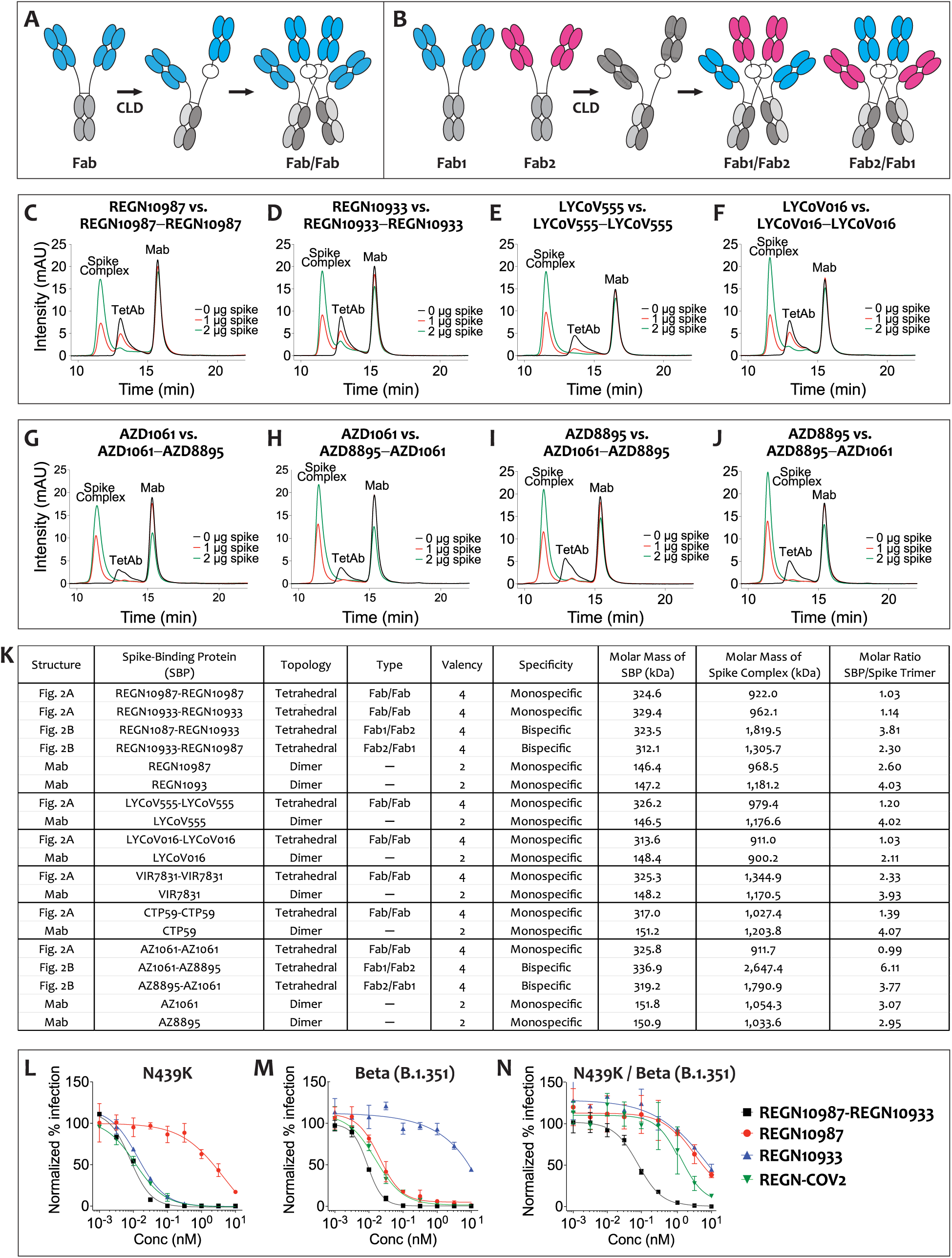
Tetrahedral formatting of Fab domains provides a distinct cooperative binding advantage over conventional monoclonal antibody planar configurations. Schematic domain configurations illustrating (**a**) a tetravalent monospecific Fab/Fab tetrahedral antibody, and (**b**) a tetravalent bispecific Fab1/Fab2 tetrahedral antibody. (**c-j**) Stoichiometric competition binding demonstrating the cooperative spike binding of Fab-Fab and Fab1-Fab2 tetrahedral antibodies monitoring complex formation with individual SARS-CoV-2 spike protein trimers across a titration range of 0 to 2 μg using (**c-f**) Fab/Fab tetrahedral antibodies mixed with parent antibody, **(c)** REGN10987-REGN10987 mixed with REGN10987, **(d)** REGN10933-REGN10933 mixed with REGN10933, **(e)** LYCoV555-LYCoV555 mixed with LYCoV555, and **(d)** LYCoV016-LYCoV016 mixed with LYCoV016, and (**g-j**) bispecific Fab1/Fab2 tetrahedral antibodies mixed with each parent antibodies, (**g**) AZD1061-AZD8895 mixed with AZD1061, (**h**) AZD8895-AZD1061 mixed with AZD1061 (**i**) AZD1061-AZD8895 mixed with AZD8895, and (**j**) AZD8895-AZD1061 mixed with AZD8895. (**k**) Stoichiometric binding complex summary table for monospecific Fab-Fab and bispecific Fab1-Fab2 tetrahedral antibodies, and parent antibodies, detailing molecular configurations, topological assignments, valency, specificity, molar mass of the spike binding protein, molar mass of the spike complex, and the calculated molar ratio of the spike binding protein to the spike trimer in the spike complex. (**l-n**) Bispecific Fab1/Fab2 tetrahedral antibodies are more potent than two-antibody cocktails. Pseudovirus neutralization demonstrating order of magnitude superior activity of the bispecific REGN10987-REGN10933 tetrahedral antibody compared with its parent antibodies REGN10987 and REGN10933, and with a two-antibody cocktail of REGN10987 and REGN10933 (REGN-CoV2), against three resistant SARS-CoV-2 mutant variants, (**l**) the N439K variant, (**m**) the Beta (B.1.351) variant, and (**n**) the N439K/Beta (B.1.351) double variant.

Pseudovirus neutralization assays confirmed that this architectural reformatting consistently translates to pronounced enhancements in antiviral activity (**Supplementary Fig. 8**). Notably, the engineered Fab-Fab tetrahedral antibodies RG2-RG2 and CT1-CT1 exhibited an approximately 100-fold increase in neutralization potency against the Beta B.1.351 variant relative to their respective parental monoclonal antibodies, REGN10933 and CT-P59 (**Supplementary Table 3**).

### Topological Requirements for Cooperative Intra-Spike Engagement

Our biophysical analyses revealed an intriguing structural phenomenon: standard bivalent ACE2 dimers, regardless of whether dimerization was mediated via the CLD, the Fc domain, or both modules, bound individual spike trimers with an apparent absence of cooperative binding. Given that the SARS-CoV-2 spike trimer displays three independent receptor-binding domains (RBDs), this limited binding suggests that the spatial layout of native RBDs is sterically incompatible with concurrent bivalent engagement by conventional planar formats. Conversely, the pronounced binding exhibited by our tetravalent ACE2-740/615, ACE2-740/Fab, Fab/Fab, and Fab1/Fab2 tetrahedral antibodies indicate that their three-dimensional, non-planar configuration allows the molecule to adapt to spatial constraints and achieve true intra-spike multivalent engagement. Characterization of the absolute molar masses of these tetrahedral-antibody–spike complexes under conditions of antibody excess (**Fig. 2k**) revealed two distinct mechanistic modalities of intra-spike interaction:

Strict 1:1 Stoichiometric Complexes. Observed for ACE2-740/615 and the majority of Fab/Fab monospecific tetrahedral antibodies. This reflects an intra-spike, inter-subunit interaction wherein a single tetrahedral molecule simultaneously bridges epitopes across at least two of the three identical S protein protomers within a single trimer.

Greater than 2:1 Stoichiometric Complexes. Exemplified by ACE2-740/B13A and all bispecific Fab1/Fab2 tetrahedral antibodies. This configuration indicates an intra-spike engagement profile where each individual tetrahedral assembly occupies distinct epitopes in a manner that leaves adjacent domains uncrowded, allowing multiple tetrahedral molecules to concurrently access a single spike trimer without inducing steric clash.

Collectively, these structural insights indicate that the tetrahedral configuration provides a robust, universal architecture for overcoming the geometric limitations that prevent conventional immunoglobulins from achieving simultaneous, multivalent target engagement.

### Broad-Spectrum Neutralization of Highly Mutable Viral Variants

A major vulnerability of traditional monoclonal antibodies targeting SARS-CoV-2 is their susceptibility to mutational escape, as evidenced by the widespread loss of clinical efficacy against highly drifted evolutionary lineages. The emergence of the heavily mutated Omicron (B.1.1.529) variant completely compromised or abolished the neutralizing activity of most historically authorized therapeutic antibodies. This clinical challenge highlights the unique structural advantage of the tetrahedral ACE2 presentation: because the virus relies on ACE2 engagement for cellular entry, it cannot easily mutate its binding interface without a catastrophic loss of biological fitness.

Both the ACE2-740/615 (designated HB1507) and ACE2-740/B13A (designated HB1516) tetrahedral antibodies demonstrated exceptional, broad-spectrum neutralization across a panel of twelve distinct viral variants, including Omicron B.1.1.529 (**Fig. 3**). Notably, the tetravalent ACE2-740/615 tetrahedral antibody neutralized Omicron B.1.1.529 with extreme potency, yielding a half-maximal inhibitory concentration (*IC₅₀*) value of 16 pM. This represents a two order-of-magnitude or greater increase in neutralization efficacy compared to a comprehensive benchmarking panel of established monoclonals, including REGN10987, REGN10933, LY-CoV555, LY-CoV016, AZD1061, AZD8895, VIR-7831, and CT-P59 (**Fig. 3**, **Supplementary Table 4**). Across all twelve lineages tested, the bispecific ACE2-740/B13A tetrahedral antibody delivered equivalent or superior neutralization profiles relative to the identical eight-antibody reference panel. These findings confirm that presenting the native receptor within a multi-copy, tetrahedral layout yields an ultra-potent, escape-resistant therapeutic framework capable of neutralizing highly transmissible, resistant variants.

**Fig. 3.**
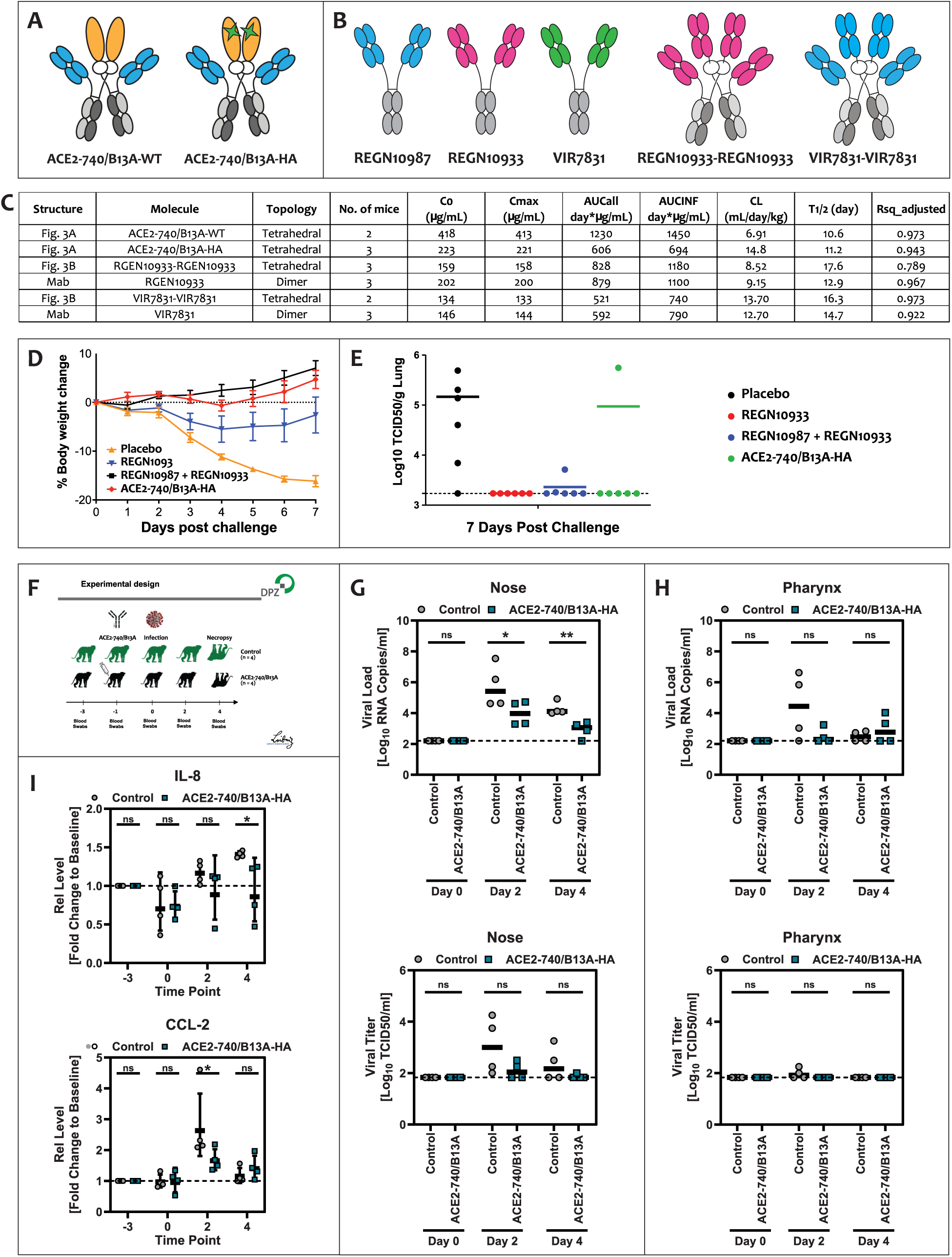
Tetrahedral antibodies demonstrate antibody-like pharmacokinetics and protection against a viral escape variant *in vivo*. Structural layouts of (**a**) the catalytically active and inactive bispecific ACE2-740/B13A candidates and (**b**) tetrahedral Fab-Fab architectures incorporating variable regions derived from the regulatory-authorized monoclonal antibodies REGN10933 and VIR-7831. (**c**) Summary table of pharmacokinetic properties following a single intravenous bolus administration (10 mg/kg) in human FcRn transgenic (Tg32) mice of bispecific ACE2-740/B13A tetrahedral antibodies, monospecific Fab-Fab tetrahedral antibodies, and parental controls, detailing molecular configurations, topological assignments, and derived pharmacokinetic parameters (C₀, C_max_, AUC_all_, AUC_INF_, CL, and t_1/2_). (**d, e**) Prophylactic efficacy of ACE2-740/B13A-HA in golden Syrian hamsters (*Mesocricetus auratus*, *n* = 6 animals per group) treated via a single intraperitoneal injection on day –1, challenged with 1.09 x 10⁵ PFU of SARS-CoV-2 (Beta B.1.351 variant) on day 0, and evaluated for (**d**) daily body weight changes across days 1–7 post-infection, and (**e**) viral titers (log₁₀ TCID₅₀/g) isolated from lung homogenates. Statistical indicators in (**d**): ACE2-740/B13A-HA vs. placebo (*p* < 0.0001); ACE2-740/B13A-HA vs. REGN10987+REGN10933 cocktail (*p* = NS); and REGN10933 vs. placebo (*p* = 0.0775). (**f**) Experimental schema for the non-human primate challenge model using rhesus macaques (*Macaca mulatta*) randomized to an untreated control group (*n* = 4) or a prophylaxis group (*n* = 4) receiving intravenous ACE2-740/B13A-HA 24 hours prior to infection with 1 x 10⁵ PFU of the Omicron subvariant XBB.1.5. (**g, h**) Viral genomic RNA copy numbers quantified via RT-qPCR and infectious viral titers determined via TCID₅₀ assays from nasal and pharyngeal swabs sampled at days 0, 2, and 4 post-infection. (**i**) Systemic IL-8 and CCL2 plasma concentrations (pg/mL) tracking inflammatory kinetics at days –3, 0, 2, and 4 post-challenge. Lines represent longitudinal measurements of individual animals. Statistical significance was modeled using a one-sided, unpaired Student’s *t*-test with Welch’s correction for (**g, h**), and two-way analysis of variance (ANOVA) with Šídák’s multiple comparisons test for (**i**).

### Mitigating Mutational Escape with Bispecific Tetrahedral Fab Formats

To determine whether structurally integrated multi-valent configurations could neutralize escape-prone viral variants more effectively than corresponding monoclonal combinations, we engineered a panel of bispecific tetrahedral Fab antibodies incorporating two distinct anti-spike variable domains. A bispecific tetrahedral Fab candidate, designated HB1701 (**Fig. 2b**), was constructed utilizing the fragment antigen-binding (Fab) sequences derived from the monoclonal antibodies REGN10987 and REGN10933.

In pseudovirus neutralization assays, HB1701 neutralized the highly resistant N439K/B.1.351 viral variant with a 25-fold potency enhancement relative to an equimolar, two-antibody cocktail of its parental monoclonal references (**Figs. 2l-n**, **Supplementary Table 5**). The absolute neutralizing capacity of HB1701 was comparable to the highly potent ACE2-740/615 and ACE2-740/B13A tetrahedral architectures. These data demonstrate that spatial formatting within a bispecific tetrahedral geometry can overcome mutational resistance profiles that render standard clinical antibody cocktails ineffective.

### Pharmacokinetics and In Vivo Prophylactic Efficacy of Antiviral Candidates

To evaluate the translational viability of these novel architectures, we determined the plasma half-lives of our tetrahedral antibodies using homozygous human neonatal Fc receptor (FcRn) Tg32 transgenic mice, a model known to predict human clearance rates with high accuracy.^17^ Following a single intravenous (i.v.) bolus administration at a dose of 10 mg/kg, functional plasma concentration profiles were quantified via spike-binding electrochemiluminescence immunoassay (ECLIA) (**Supplementary Figs. 9, 10**).

Two distinct versions of the ACE2-740/B13A tetrahedral antibody, one maintaining native catalytic angiotensin-converting enzyme activity (ACE2-740/B13A-WT) and one rendered catalytically inactive via targeted active-site mutation (ACE2-740/B13A-HA) (**Supplementary Fig. 11**, **Supplementary Table 6**), demonstrated prolonged terminal half-lives (t_1/2_) of 10.6 and 11.2 days, respectively. Crucially, this represents a 33.5-fold and 35.4-fold extension in plasma residence time compared to the clinical terminal half-life reported for soluble recombinant human monomeric ACE2 (APN01) in humans (t_1/2_ = 7.6 hours following a 1.2 mg/kg i.v. bolus).^18^

Furthermore, two pure antibody tetrahedral variants, designated REGN10933-REGN10933 and VIR7831-VIR7831, which incorporate the variable domains of REGN10933 and VIR-7831, respectively, exhibited terminal half-lives of 17.6 and 16.3 days. These pharmacokinetic profiles compare favorably with the typical clearance parameters observed for the full-length REGN10933 and VIR7831 parental immunoglobulin G (IgG) molecules (12.9 and 14.7 days, respectively). These data demonstrate that the integration of the collectrin-like domain (CLD) tetrahedral nucleus does not accelerate circulatory clearance or destabilize endosomal FcRn-mediated recycling mechanisms.

We next translated these in vitro findings into a rigorous in vivo prophylactic antiviral model of SARS-CoV-2 infection in golden Syrian hamsters (*Mesocricetus auratus*).^19^ Animal cohorts treated with a single prophylactic dose of the ACE2-740/B13A-HA tetrahedral antibody (25 mg/kg) via intraperitoneal injection were completely protected against infection-induced weight loss. This protective efficacy was comparable to a high-dose clinical cocktail consisting of two distinct monoclonal antibodies, REGN10987 (25 mg/kg) and REGN10933 (25 mg/kg), and significantly superior to monotherapy with REGN10933 alone at an equivalent dose (25 mg/kg; **Figs. 3d, e**).

Additionally, the in vivo protective efficacy of ACE2-740/B13A-HA was validated in a non-human primate challenge model using rhesus macaques (*Macaca mulatta*). Four macaques per cohort were left untreated or received the tetrahedral compound intravenously at 24 hours prior to intranasal and intratracheal challenge with the Omicron subvariant XBB.1.5 (**Fig. 3f**). The animals were monitored for clinical signs, viral load (genomic RNA), viral titer (median tissue culture infectious dose; TCID₅₀) in nasal and pharyngeal swabs, and systemic cytokine levels in plasma. No overt clinical signs of disease were observed post-challenge.

Viral RNA was readily detectable in the nose (**Fig. 3g**) and to a lesser degree in the pharynx (**Fig. 3h**) using quantitative reverse transcription PCR (RT-qPCR) for viral genomic RNA detection. Prophylaxis with ACE2-740/B13A-HA significantly decreased viral RNA copy numbers in the nasal epithelium at 2 and 4 days post-infection. In the pharynx, ACE2-740/B13A-HA prophylaxis also reduced viral load at 2 days post-infection; by day 4 post-infection, viral loads in both control and treated animals had declined to background levels.

Concordant results were obtained when infectious virus was quantified using the TCID₅₀ viral titer assay. Finally, profiling of the systemic immune response revealed that ACE2-740/B13A-HA prophylaxis reduced C-C motif chemokine ligand 2 (CCL2) and interleukin-8 (IL-8) levels in the plasma of infected animals (**Fig. 3i**), in agreement with clinical data demonstrating upregulation of these inflammatory factors in COVID-19 patients and SARS-CoV-2 infected macaques [21– 25]. At 4 days post-infection, treated animals also showed increased interferon-beta (IFN-beta) and tumor necrosis factor-alpha (TNF-alpha) levels relative to untreated controls (**Supplementary Fig. 12**), suggesting a shift in the quality and kinetics of the antiviral innate immune response.

Consistent with this favorable translational profile, repeated intravenous bolus safety evaluation of ACE2-740/B13A in cynomolgus monkeys (*Macaca fascicularis*) at doses up to 100 mg/kg/day administered twice weekly for 4 weeks produced no treatment-related adverse findings, establishing a no-observed-adverse-effect level (NOAEL) of 100 mg/kg/day. Collectively, these non-human primate data demonstrate that the tetrahedral antibody framework significantly suppresses both viral replication and pathogenic cytokine expression in vivo.

### Dual-Targeting of Soluble Cytokines and Blending Inflammatory Synergy

We next examined whether the tetrahedral architecture could be applied to capture soluble signaling proteins, specifically targeting synergistic cytokines that drive chronic inflammatory and autoimmune pathologies.^20^ We generated bispecific tetrahedral Fab antibodies (**Fig. 4a**) using the variable domains of adalimumab (anti-tumor necrosis factor-alpha; anti-TNF-alpha) and secukinumab (anti-interleukin-17A; anti-IL-17A).

**Fig. 4.**
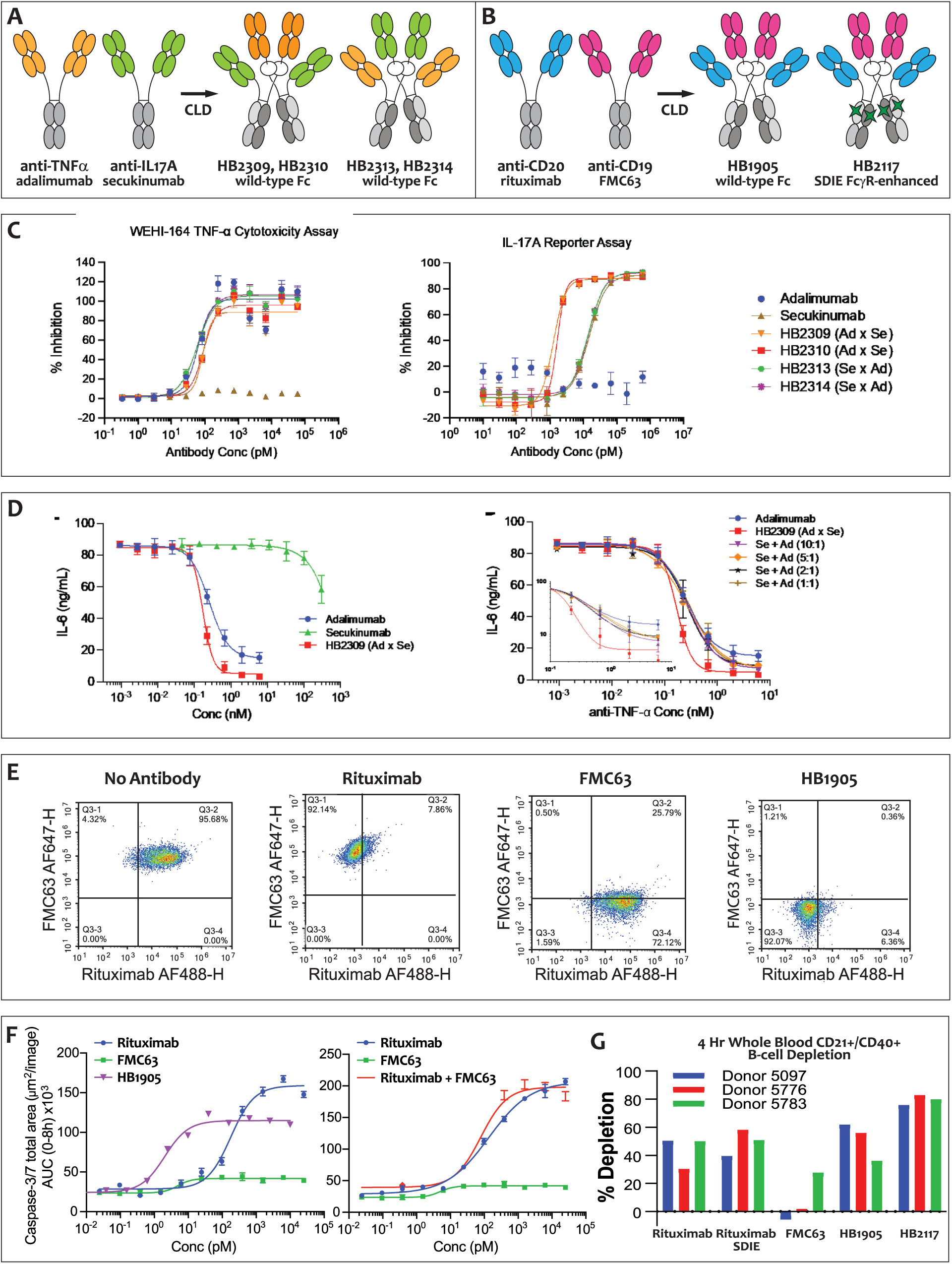
Simultaneous multivalent targeting of TNFα and IL17A drives anti-cytokine synergy; simultaneous multivalent targeting of CD19, CD20, and low-affinity Fc*γ* receptors drives enhanced effector cell recruitment and B-cell depletion. (**a**) Schematic domain layout of anti-TNF-α/anti-IL-17A bispecific Fab1/Fab2 tetrahedral antibodies. (**b**) Schematic domain layout of dual-Fc bispecific anti-CD19/anti-CD20 tetrahedral antibodies comparing the wild-type Fc variant HB1905 with the affinity-enhanced, symmetrically engineered SDIE variants HB2117. (**c**) Neutralizing activity of anti-TNF-α/anti-IL-17A bispecific Fab1/Fab2 tetrahedral antibodies compared against their parent antibodies adalimumab and secukinumab. Neutralization of TNF-α induced cytotoxicity in WEHI-164 cells by tetrahedral antibodies HB2309 and HB2310 (Ad x Se configuration), and HB2313 and HB2314 (Se x Ad configuration) compared with the adalimumab and secukinumab parent antibodies (left panel), and neutralization of IL-17A activity in an IL-17A reporter assay by HB2309 and HB2310 (Ad x Se) and HB2313 and HB2314 (Se x Ad) compared with the adalimumab and secukinumab parent antibodies (right panel). (**d**) Neutralization of TNF-α/IL-17A induction of IL-6 secretion by normal human dermal fibroblasts (NHDF) by the Ad x Se tetrahedral antibody (HB2309), compared to the secukinumab and adalimumab parent antibodies (left panel). The HB2309 Ad x Se tetrahedral antibody compared to adalimumab and several two-antibody cocktails consisting of the secukinumab and adalimumab parent antibodies at ratios of 1:1, 2:1, 5:1, and 10:1 (right panel). The x-axis reflects the concentration of the anti-TNF-α Fab portion that is present in the HB2309 Ad x Se tetrahedral antibody, the adalimumab control, and in each of the two-antibody cocktails. The inset shows an expanded view of the dose response plotted in log-log scale. (**e**) Binding analysis of anti-CD20/anti-CD19 bispecific Fab1/Fab2 tetrahedral antibodies for binding of cell surface CD20 and CD19 compared against their parent antibodies (rituximab, FMC63). Flow cytometry analysis of CD20/CD19-positive Toledo cells labelled by incubation with AF488-labeled rituximab and AF647-labeled FMC63 following pre-incubation of cells with no antibody, unlabeled rituximab, unlabeled FMC63, and tetrahedral antibody HB1905. (**f**) Kinetic activation profiles of fluorogenic caspase-3/7 cleavage tracking antibody-dependent cellular cytotoxicity (ADCC) in co-cultures of Toledo target B cells and primary human NK effector cells (E:T ratio = 10:1), demonstrating significantly enhanced apoptotic potency for HB1905 (left panel) compared to an equimolar combination cocktail of monoclonal rituximab and FMC63 references (right panel) relative to baseline standalone parental monomers. (**g**) Ex vivo human whole-blood B-cell depletion profiles showing that the SDIE-enhanced tetrahedral variant HB2117 mediates higher maximal target cell depletion than HB1905, rituximab, FMC63, or an isolated SDIE-enhanced rituximab control across independent healthy donors.

To evaluate the impact of spatial configuration on soluble ligand capture, we expressed two alternative tetrahedral layouts: one featuring the adalimumab Fab fused to the collectrin-like domain (CLD) core with the secukinumab Fab linked to the crystallizable fragment (Fc) domain (Ad x Se), and a reciprocal variant featuring the opposite orientation (Se x Ad). Both tetrahedral configurations neutralized recombinant human TNF-alpha with functional potencies comparable to native, monoclonal adalimumab (**Fig. 4c**, **Supplementary Table 7**).

By contrast, the spatial orientation heavily dictated target engagement kinetics for the secondary cytokine; the Ad x Se tetrahedral configuration neutralized human IL-17A with a potently enhanced 8.3-fold to 11.5-fold advantage over the reciprocal Se x Ad layout, and a 9.1-fold to 12.0-fold potency improvement over monoclonal secukinumab itself (**Fig. 4c**, **Supplementary Table 7**). Based on these optimization parameters, the Ad x Se tetrahedral variant, designated HB2309, was selected to evaluate the inhibition of cytokine-mediated inflammatory synergy using normal human dermal fibroblasts (NHDF). Simultaneous exposure to TNF-alpha and IL-17A stimulates a hyper-inflammatory response in NHDF populations, driving significantly elevated secretion of interleukin-6 (IL-6) compared to stimulation by either isolated cytokine alone (**Supplementary Fig. 13**).

As demonstrated in **Fig. 4d**, HB2309 abrogated the synergistic induction of IL-6 mediated by the TNF-alpha and IL-17A combination, a therapeutic endpoint that neither standalone adalimumab nor standalone secukinumab could achieve. Furthermore, HB2309 fully suppressed the inflammatory IL-6 cascade an order of magnitude more effectively than a comprehensive benchmarking panel of traditional two-antibody combinations of secukinumab and adalimumab blended across molar ratios ranging from 1:1 to 10:1 (**Fig. 4d**). These findings establish that the modular tetrahedral framework can be structurally optimized to systematically disrupt multi-cytokine inflammatory networks.

### Simultaneous Bivalent Targeting of CD19, CD20, and Fc*γ* Receptors for B-cell Depletion

To evaluate tetrahedral antibodies as natural killer (NK) cell and monocyte engagers, we designed molecules capable of simultaneous dual engagement of B-cell surface antigens and Fcγ receptors (FcγRs). CD19 and CD20 were selected as targets for B-cell depletion in malignancies and autoimmune disorders to mitigate lineage escape and tumor recurrence. While co-targeting these markers is a validated chimeric antigen receptor (CAR)-T cell strategy,^21–23^ an effective, off-the-shelf alternative remains an unfulfilled clinical need.

We generated two dual-Fc anti-CD20/anti-CD19 bispecific tetrahedral antibodies, designated HB1905 and HB2117, featuring wild-type and SDIE-enhanced (Ser239Asp/Ile332Glu) dual Fc domains, respectively. These molecules incorporate Fab domains from the chimeric antibodies rituximab and FMC63.^24,25^ Surface plasmon resonance (SPR) analyses verified that monovalent binding affinities to CD20, CD19, FcγRs, and complement component C1q were comparable to those of the parental antibodies (**Supplementary Fig. 14**) **[26]**. Pre-incubation of HB1905 with CD20/CD19-double-positive human Toledo B cells completely occluded subsequent binding of fluorescently labeled rituximab and FMC63 (**Fig. 4e**), confirming simultaneous steric target engagement.

To assess functional cytotoxicity, we quantified apoptosis as an indicator of antibody-dependent cellular cytotoxicity (ADCC) using Toledo cells co-cultured with primary human NK cells. HB1905 demonstrated a 39-fold increase in cytotoxic potency relative to a combination cocktail of rituximab and FMC63 (**Fig. 4f**, **Supplementary Table 8**). Crucially, HB2117 mediated robust B-cell depletion in ex vivo human whole-blood assays across three independent healthy donors, achieving higher maximal B-cell ablation than HB1905, rituximab, FMC63, or SDIE-enhanced rituximab (**Fig. 4g**). These results indicate that the dual-Fc SDIE-enhanced configuration represents a new standard in B-cell depletion.

To clarify the translational relevance of this platform, we engineered HB2174, a humanized version of HB1905, and HB2182 and HB2190, humanized versions of HB2117, but differing by incorporating asymmetric instead of symmetric SDIE mutations (**Fig. 5a**). We then profiled binding against recombinant Chinese hamster ovary (CHO) cells expressing individual FcγRs. We observed superior binding across all six low-affinity FcγRs tested (FcγRIIIa-158F, FcγRIIIa-158V, FcγRIIIb, FcγRIIa-131H, FcγRIIb, and FcγRIIc) compared to antibodies with wild-type Fc domains (rituximab and huFMC63), and SDIE-enhanced variants (rituximab-SDIE, FMC63-SDIE, and tafasitamab). Notably, binding of all three humanized constructs to FcγRIIIa, the primary activating receptor on NK cells and monocytes, was up to two orders of magnitude greater than that of rituximab and huFMC63, and up to one order of magnitude superior to that of rituximab-SDIE, FMC63-SDIE, and tafasitamab (**Fig. 5b, c**). Contrasting these multivalent cell-binding profiles with monovalent affinity constants (*K_D_*) derived via Carterra surface plasmon resonance (SPR) confirmed that superior cell-surface binding observed for the dual Fc tetrahedral antibodies reflects functional cooperativity of the respective dual-Fc domain architectures toward low-affinity FcγRs (**Fig. 5d**).

**Fig. 5.**
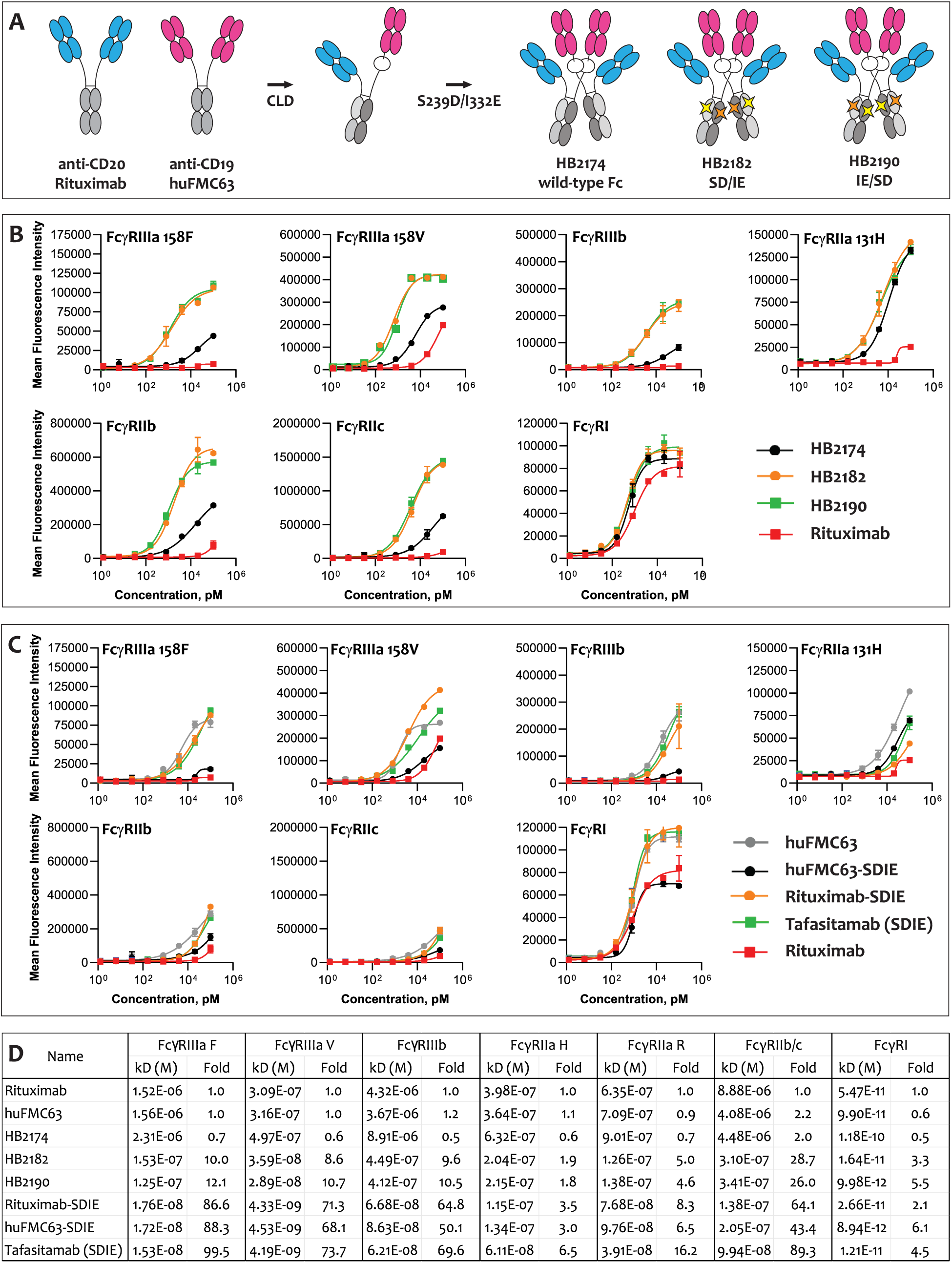
Fc domain engineering tunes cell-surface binding cooperativity. (**a**) Schematic architectures of dual-Fc anti-CD19/anti-CD20 tetrahedral antibodies engineered with wild-type, asymmetric SD/IE-enhanced, and asymmetric IE/SD-enhanced dual Fc domains. (**b, c**) Equilibrium binding titrations evaluated against recombinant CHO cells stably expressing individual human low-affinity Fc receptors, including FcγRIIIa-158F, FcγRIIIa-158V, FcγRIIIb, FcγRIIa-131H, FcγRIIb, and FcγRIIc, demonstrating functional cooperativity for wild-type and enhanced dual Fc domain architectures compared to (**b**) rituximab and (**c**) wild-type rituximab, huFMC63 and SDIE-enhanced benchmarks (rituximab-SDIE, huFMC63-SDIE, tafasitamab). (**d**) Monovalent baseline binding affinities determined by surface plasmon resonance (SPR) for the tetrahedral antibodies and antibody controls shown in (**b**) and (**c**). Cross-referencing these SPR values with the CHO-cell binding profiles confirms that multivalent functional cooperativity successfully overcompensates for reduced monovalent affinity. This cooperative effect is driven by both the wild-type and asymmetric dual-Fc tetrahedral antibody configurations.

Collectively, these data demonstrate that the physical integration of both dual-targeting Fc and Fab domains into a tetrahedral architecture substantially enhances effector cell recruitment and B-cell depletion compared to established antibody therapies. Based on these and other recently published findings,^26^ a candidate has progressed into human Phase I clinical trials evaluating safety and tolerability in healthy volunteers and participants with moderately to severely active systemic lupus erythematosus (SLE).^27,28^

### Engineering Covalent Stability via CLD Interface Disulfide Bonds

To optimize the structural stability of the non-native tetrahedral symmetry, we introduced targeted, disulfide-forming cysteine mutations into the CLD dimerization interface. The dimer interface formed by the pairing of the first and second CLD polypeptides (**Fig. 6a**) is mediated by extensive polar interactions across the second (residues 636 to 658) and fourth (residues 708 to 717) helices of each CLD chain (positions mapped relative to human ACE2, UniProt Q9BYF1). Within a dual-Fc bispecific anti-CD19/CD20 tetrahedral antibody framework, we evaluated two rational cysteine substitutions, Ala714Cys within the fourth helix (**Fig. 6b**) and Ser645Cys within the second helix (**Fig. 6c**), as well as a Ser645Cys/Ala714Cys double substitution.

**Fig. 6.**
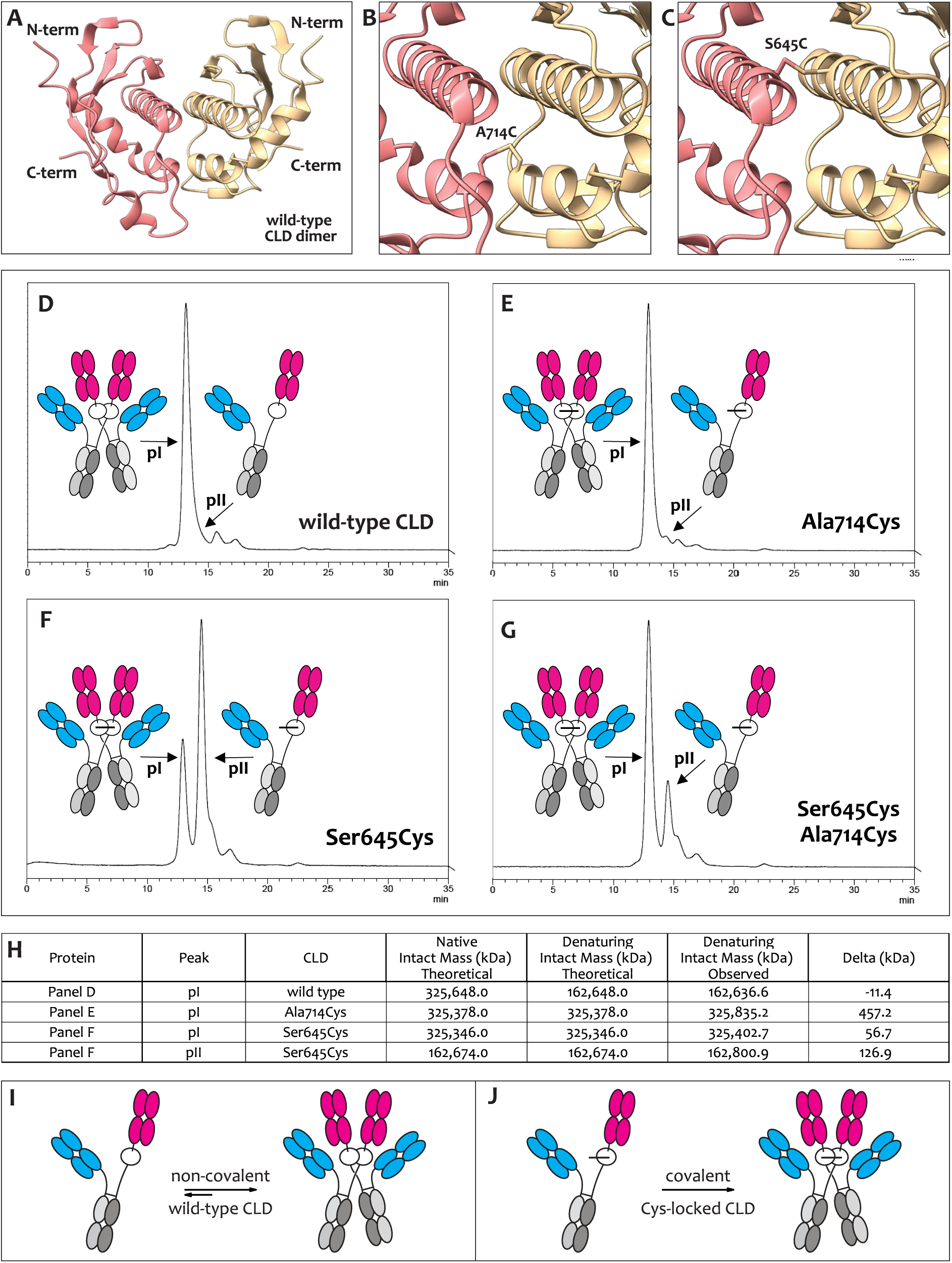
Interface disulfide engineering covalently locks the quaternary tetrahedral topology. (**a**) Cryo-EM structural density mapping of the wild-type collectrin-like domain (CLD) dimer (PDB ID: 6M18) highlighting the spatial orientation of the four projecting amino and carboxy termini that direct tetrahedral antibody self-assembly. (**b, c**) Structure-guided computational molecular modeling illustrating the introduction of targeted, interface disulfide-bond-forming mutations, comparing (**b**) the fourth-helix Ala714Cys substitution against (**c**) the second-helix Ser645Cys substitution. (**d–f**) Analytical size-exclusion chromatography (SE-HPLC) profiles of bispecific anti-CD20/CD19 tetrahedral variants under high flow and high pressure, demonstrating concentration-dependent dissociation-reassociation tailing equilibria for (**d**) the non-covalent wild-type CLD baseline, complete covalent locking and peak symmetry for (**e**) the fourth-helix Ala714Cys variant, and severe structural disruption or incomplete quaternary self-assembly for (**f**) the second-helix Ser645Cys variant, and (**g**) the Ser645Cys Ala714Cys double variant. (**h**) Denaturing intact mass spectrometry results for the major species observed in panels **d-f**. (**i, j**) Summary of the equilibrium behavior observed for tetrahedral antibodies with CLD dimers held together by (**i**) strong non-covalent forces or **(j**) the Ala714Cys mutation.

Analytical SE-HPLC demonstrated that the Ala714Cys substitution successfully locked the tetrahedral architecture. Chromatographic integration revealed that the non-covalent wild-type tetrahedral antibody consisted of a 94.27% main peak fraction (pI) and a 5.73% product-related impurity trail, which exhibited prominent peak asymmetry ("tailing") under high pressure and flow rates. This severe asymmetry indicates a concentration-dependent dissociation-reassociation equilibrium (Fig. 6d). Using an automated line trace at an exactly normalized 6.00% fractional peak height threshold (A_280_ intensity level = 2,064), the near-baseline asymmetry factor (A_s_) was calculated at 2.38 for the non-covalent tetrahedral antibody. Conversely, the engineered Ala714Cys variant achieved a highly compact profile consisting of a 95.14% primary product fraction alongside a 4.86% product-related impurity that successfully resolved as a distinct, baseline-separated secondary peak downstream (pII; Fig. 6e). Evaluated at the identical 6.00% fractional height boundary (A_280_ intensity level = 2,309), the Cys-locked tetrahedral antibody demonstrated a significantly reduced, stabilized near-baseline A_s_ of 2.02. A minor adjacent peak (<2%) corresponded to the retention time of the non-covalent tail, representing the minimal unbonded, half-tetrahedral antibody subpopulation.

Denaturing intact mass spectrometry (MS) following near-complete de-glycosylation confirmed quantitative inter-chain disulfide bond formation of the Ala714Cys variant, yielding an observed molecular mass of 325,835.2 Da, closely matching the theoretical molecular mass of 325,378.0 Da for the fully assembled tetrahedral antibody (**Fig. 6h**). By contrast, the non-covalent, wild-type tetrahedral reference yielded an observed mass of 162,636.6 Da under identical denaturing conditions, confirming full dissociation into the constituent half-tetrahedral antibody intermediate (theoretical mass 162,648.0 Da). In summary, while tetrahedral antibodies held together solely by strong non-covalent forces are maintained as superdimers with an equilibrium strongly favoring formation (**Fig. 6i**), Ala714Cys disulfide-locked tetrahedral antibodies, as expected, do not appear to dissociate (**Fig. 6j**).

In contrast to the fourth-helix modification, SE-HPLC of the second-helix Ser645Cys variant revealed an incomplete structural assembly profile (**Fig. 6f**). This species resolved into a minor peak corresponding to the fully covalent tetrahedral assembly (observed mass: 325,402.7 Da, theoretical mass: 325,346.0 Da) and a predominant major peak matching a half-tetrahedral monomeric intermediate (observed mass: 162,800.9 Da, theoretical mass: 162,674.0 Da). These data indicate that while a minor fraction of the Ser645Cys variant achieved covalent stabilization, the mutation sterically or electrostatically disrupted the underlying non-covalent association of the remaining chains, hindering efficient self-assembly into the quaternary tetrahedral topology. The Ser645Cys/Ala714Cys dual substitution behaved similarly to Ser645Cys, indicating that disruption of dimer formation by Ser645Cys was dominant over disulfide-locking mediated by Ala714Cys (**Fig. 6g**).

## Discussion

We have demonstrated a robust and versatile platform for engineering tetrahedral antibodies, a distinct class of multispecific, multivalent therapeutics. This architecture can be universally applied to any combination of Fab fragments, Fc domains, and extracellular domains. By integrating the ACE2-derived collectrin-like domain (CLD) directly into the antibody hinge region, we engineered a non-planar, three-dimensional topology that coordinates up to six unobstructed, fully functional binding faces. This structural approach systematically overcomes the spatial constraints and localized steric hindrance that frequently limit conventional, linear or planar IgG-based multispecific formats.^3,4^ Our biophysical analyses demonstrate that this tetrahedral configuration provides a pronounced cooperative binding advantage over traditional architectures. Standard dimeric ACE2-Fc and monoclonal antibody (mAb) formats are often restricted to monovalent target engagement due to strict geometric and steric mismatches with oligomeric receptors. By contrast, the non-planar orientation of tetrahedral antibodies facilitates simultaneous, multi-site engagement of complex macro-structures, such as the trimeric SARS-CoV-2 spike protein. This multivalent engagement results in an increase of up to two orders of magnitude in functional spike-binding affinity driven by strong functional cooperativity, translating directly into the potent neutralization of highly transmissible, escape-prone viral variants, including the heavily mutated Omicron B.1.1.529 lineage.

Treatment with HB1516 (ACE2-740/B13A) significantly reduced both nasal viral load and proinflammatory cytokine expression, demonstrating protection in rhesus macaques against a highly immune-evasive Omicron lineage. These findings are consistent with those of Urano et al.,^30^ who demonstrated that administration of an engineered ACE2 decoy reduced viral replication and pulmonary disease in SARS-CoV-2 Delta-infected macaques,^29^ supporting the delivery of ACE2-based decoys as a broadly active strategy against antigenically divergent variants. Although the standalone neutralizing potency of the B13A monoclonal antibody, as with other clinical mAbs with emergency use authorization, was substantially diminished by the rapid antigenic evolution of B.1.1.529 sublineages, its physical integration within an ACE2 decoy preserves broad-spectrum activity by targeting the highly conserved structural requirement for spike glycoprotein engagement of the host receptor. This dual-targeting framework may therefore extend the effective therapeutic lifespan of an otherwise escaped monoclonal antibody while systematically increasing the genetic barrier to viral resistance. Importantly, although HB1516-treated macaques exhibited an elevation in localized IFN-beta and TNF-alpha levels at day 4 post-infection, this shift occurred strictly within the context of a heavily reduced viral burden and suppressed CCL2 and IL-8 levels. This precise signaling signature suggests an accelerated, protective antiviral innate immune response rather than generalized, treatment-induced hyper-inflammation.

Beyond infectious disease indications, the tetrahedral platform is effective at co-targeting cell-surface receptors and soluble cytokines. In B-cell lymphoma models, our bispecific anti-CD20/anti-CD19 tetrahedral antibodies exhibited a 39-fold increase in cytotoxic potency relative to corresponding parental antibody cocktails. This substantial enhancement highlights their potential to overcome established resistance mechanisms in oncology and effect an autoimmune reset in severe autoimmune diseases, such as systemic lupus erythematosus (SLE). Similarly, our bispecific anti-tumor necrosis factor-alpha (TNFα)/anti-interleukin-17A (IL-17A) tetrahedral candidate completely abolished synergistic interleukin-6 (IL-6) induction in primary human dermal fibroblasts, outperforming parental antibody cocktails by more than an order of magnitude. These functional improvements depend on precise domain orientation, demonstrating that the platform’s architectural modularity and non-planar spatial distribution of binding moieties are crucial for disrupting multi-component extracellular signaling networks.

Crucially, these tetrapartite antibodies retain favorable, traditional IgG-like pharmacological properties. They exhibit robust pharmacokinetics, with terminal elimination half-lives t_1/2_ in human neonatal Fc receptor (FcRn) transgenic mice that are comparable to those of standard parental IgG controls. These data indicate that the multivalent, macromolecular assemblies undergo normal, unhindered endosomal FcRn-mediated salvage and recycling pathways.

We have advanced the tetrahedral multispecific platform in two critical ways, significantly improving its translational relevance and structural integrity. First, humanized anti-CD19/CD20 tetrahedral antibodies (HB2174, HB2182, HB2190) exhibited superior binding across all six low-affinity FcγRs compared to conventional and SDIE-enhanced variants, with up to two orders of magnitude higher binding to FcγRIIIa. This enhanced performance is driven by functional cooperativity across the dual-Fc domain architectures, as demonstrated by comparing cell-surface binding with monovalent affinities derived from surface plasmon resonance.

Second, we successfully addressed the tetrahedral architecture’s structural stability by introducing a specific Ala714Cys substitution within the CLD dimerization interface. This engineered fourth-helix lock abolished concentration-dependent dynamic dissociation and established a stable, covalent quaternary architecture, validated by analytical SE-HPLC and denaturing mass spectrometry. Conversely, second-helix Ser645Cys modifications disrupted underlying non-covalent interactions and hindered proper self-assembly, highlighting the precision required to stabilize this robust, translatable multispecific platform.

In conclusion, our tetrahedral antibody architecture establishes a new structural paradigm that moves completely beyond the strict geometric limitations of the native Y-shaped antibody topology. By enabling simultaneous, multi-dimensional cooperative binding of complex antigens and immune effector cell receptors while maintaining favorable mammalian expression yields and classic antibody pharmacological recycling, this modular, single-species architecture provides a promising framework for the development of next-generation multispecific therapeutics.

## Acknowledgements

We thank Carin M. Rollins and Miriam L. Siekevitz, co-founders of Hinge Bio, Inc., for many valuable contributions to this work, H. Michael Shepard (Ruijin Hospital, Shanghai JiaoTong University School of Medicine) and Eric Delwart (Vitalant Research Institute) for helpful discussions; Lieza Danan (LiVeritas Biosciences) for mass spectrometry, Robert Hershberg (Hinge Bio, Inc.) for advice on protein scale-up; Catherine Lucas (Hinge Bio, Inc.) for advice on assay development and animal studies; Greg Christianson and Elena Gil (Jackson Laboratory) for pharmacokinetic studies, and Jerry Moore (Pacific BioDevelopment LLC) for PK analysis.

## Author contributions

LAT, MEF, NLSC, UE, BF, BZC, JJ, and RM designed and executed experiments, and analyzed and interpreted results. NK and RV carried out animal experimentation. CR and BR carried out RT-PCR, TCID50, animal work, and cytokine analysis. VC and CD provided virus stock for animal experimentation. SP and GS provided funding acquisition, administration, and supervision. MLG contributed critical insights into biological mechanisms and the writing of the manuscript. DJC conceived the study and compositions of matter, designed the foundational methodology, provided scientific supervision, assisted with experiments and authored the original and final drafts of the manuscript. All authors read and approved the final manuscript.

## Declarations

### Ethics approval and consent to participate

All methods were carried out in accordance with the relevant local guidelines and regulations, and all experimental protocols were approved by the local institutional committees. Pharmacokinetic studies in mice were carried out at the Jackson Laboratory (JAX Services, Bar Harbor, ME, USA or JAX Services, Gainesville, FL, USA). The experiments and procedures used were approved by the Animal Care and Use Committee at The Jackson Laboratory and performed in accordance with the OLAW guidelines and were reviewed and approved by the Institutional Animal Care and Use Committee at The Jackson Laboratory. Hamster studies were carried out at BIOQUAL, Inc. (Rockville, MD, USA). Housing and handling of the animals was performed in accordance with the standards of the AAALAC International’s reference resource and the 2015 reprint of the Public Health Service (PHS) Policy on Human Care and Use of Laboratory Animals. Handling of specimens and animals was performed in compliance with the Biosafety in Microbiological and Biomedical Laboratories (BMBL). This study was performed under an IACUC-approved Animal Care and Use Protocol (attached). Rhesus (Macaca mulatta) studies were carried out at the German Primate Center–Leibniz Institute for Primate Research (Göttingen, Germany). Animals were kept in compliance with the regulations of the European Parliament and the Council Directive on the protection of animals used for scientific purposes (2010/63/EU). The experiment was approved by the Lower Saxony State Office for Consumer Protection and Food Safety (approval no.: 33.19-42502-04-20/3601). No human participants were directly involved with the study, while human samples were commercially purchased for in vitro experiments. Human healthy whole blood was obtained from Stanford Blood Center, with informed consent and approvals by Institutional Review Boards of Stanford Blood Center.

## Additional information

Correspondence should be addressed to DJC.

## Methods

### Mammalian Expression and Purification of Recombinant Proteins

Recombinant tetrahedral antibodies and associated reference proteins were expressed via transient transfection in CHO-K1 cells adapted to serum-free suspension culture (TunaCHO; LakePharma). Cells were expanded in suspension using a chemically defined, serum-free medium in shake flasks. On the day of transfection, cells were seeded into a fresh vessel at high density, and transient transfections were executed by introducing transfection reagents complexed with plasmid DNA encoding the respective heavy and light chain sequences. Post-transfection, cultures were maintained as batch-fed suspension cultures in shake-flasks. The conditioned cell culture fluid was harvested after 7 to 14 days of production, clarified by centrifugation, and passed through a 0.22 micrometer sterile filtration membrane prior to chromatographic isolation.

Affinity chromatography was performed by applying the clarified culture supernatant directly onto a column packed with CaptivA Protein A Affinity Resin (Repligen) pre-equilibrated with phosphate-buffered saline (PBS; 137 mM NaCl, 2.7 mM KCl, 10 mM Na2HPO4, 2 mM KH2PO4, pH 7.4). The column was washed with PBS until the optical density at 280 nm (OD280) returned to baseline. Target proteins were eluted with 0.25% acetic acid at pH 3.5. Eluted fractions were immediately neutralized by collecting directly into tubes containing 1 M HEPES buffer. Fractions containing the target protein were pooled, buffer-exchanged into a storage formulation of 100 mM HEPES, 100 mM NaCl, 50 mM sodium acetate, pH 6.0, filtered through a 0.2 micrometer membrane filter, and stored at 4 degrees Celsius. Final protein concentrations were calculated based on the measured OD280 value and the respective sequence-derived extinction coefficient.

## Preparative Size-Exclusion Chromatography

Preparative size-exclusion chromatography (SEC) was performed on an AKTA Avant 25 system (Cytiva). Purified protein pools were concentrated to a target range of 10–15 mg/mL using an Amicon Ultra-15 centrifugal filter unit with a 3,000 molecular weight cut-off (3MWCO; Millipore, Cat#UFC900324). Concentrated samples were loaded onto a HiLoad 26/600 Superdex 200 preparative grade column (Cytiva, Cat#28989336). Isocratic elution was conducted with a mobile phase of PBS (UCSF Cell Culture Facility, Cat#CCFAL003). Fractions corresponding to the target tetrahedral antibody assembly were identified by analytical SEC and confirmed by both reducing and non-reducing sodium dodecyl sulfate-polyacrylamide gel electrophoresis (SDS-PAGE). The validated target fractions were pooled, concentrated via ultrafiltration, and stored at 4 degrees Celsius.

## Analytical Size-Exclusion High-Performance Liquid Chromatography

Analytical size-exclusion high-performance liquid chromatography (SE-HPLC) was performed using a Prominence HPLC system (Shimadzu). Two Zenix-C SEC-300 columns (3 micrometer particle size, 300 angstrom pore size, 4.6 x 300 mm; Sepax, Cat#233300-4630) were operated in series. The mobile phase consisted of PBS supplemented with 300 mM NaCl, delivered at a constant flow rate of 0.2 mL/min. The column temperature was maintained at 25 degrees Celsius, and eluting peaks were detected by ultraviolet (UV) absorbance at 280 nm and 214 nm. Chromatographic data acquisition and peak integration were managed using LabSolutions v5.9 software (Shimadzu).

## Multi-Angle Light Scattering Analysis

The absolute molar mass and polydispersity of the engineered protein assemblies were determined by coupled inline SE-HPLC and multi-angle light scattering (SEC-MALS). The analytical system comprises the Prominence HPLC fluidics configured in series with an Optilab T-rEX refractive index (RI) detector and a Treos MALS detector (Wyatt Technology). The operating temperature for both detectors was maintained at 25 degrees Celsius. Detectors were normalized and corrected for inter-detector band broadening using the signal generated by a monomeric bovine serum albumin (BSA) reference standard. Light scattering intensity and refractive index data were acquired and processed using Astra v6.1.7 software (Wyatt Technology). A specific refractive index increment (dn/dc) value of 0.185 mL/g was used for all absolute molecular mass calculations.

## IdeZ Cleavage Analysis and TCEP Reduction

Enzymatic fragmentation was performed by incubating 100 micrograms of purified tetrahedral antibody with 3 microliters of IdeZ Protease (New England Biolabs, Cat#P0770S) in a total reaction volume of 100 microliters containing 1X GlycoBuffer 2 at 37 degrees Celsius for 60 min. To isolate fragments lacking the Fc domains, the reaction mixture was incubated with a 100 microliter slurry of Protein A Sepharose Fast Flow beads (Cytiva, Cat#17507901) for 1 hour at room temperature under constant vertical rotation. The unbound fractions containing the non-Fc fragments were recovered by centrifugation at 1,000g for 2 min using 0.5 mL spin columns (Thermo Fisher, Cat#69705). Residual protein was recovered by washing the resin with 25 microliters of PBS, and the eluates were pooled. One half of the isolated pool was reduced by adding tris(2-carboxyethyl)phosphine hydrochloride (TCEP-HCl; Thermo Fisher, Cat#20490) to a final concentration of 0.17 mM, followed by incubation for 30 min in the dark at room temperature. Both the intact IdeZ-digested and the IdeZ-digested, TCEP-reduced samples were immediately analyzed by SEC-MALS.

## Stoichiometric Competition Binding Analysis

Binding stoichiometries were evaluated by titrating the tetrahedral spike-binding candidate against aggregate-free, individual recombinant SARS-CoV-2 spike trimers (Wuhan-Hu-1 variant; GenBank: MN908947, NCBI reference sequence: NC_045512.2; ExcellGene). Binding reactions were prepared in a PBS matrix, incubated at 14 degrees Celsius for 3 hours to achieve equilibrium, and subsequently resolved and analyzed by inline SEC-MALS.

## Kinetic Exclusion Assay

Equilibrium binding affinities were determined at room temperature (approximately 22 degrees Celsius) using a Kinetic Exclusion Assay (KinExA) 3200 platform (Sapidyne Instruments). The tetrahedral spike-binding protein was held at a constant concentration to act as the constant binding partner (CBP) and titrated against serial dilutions of the recombinant spike trimer (ExcellGene) acting as the titrant. Assays were conducted in a running buffer of PBS supplemented with 1 mg/mL BSA (Sapidyne, Cat#2A3018). Following equilibrium incubation, the remaining unbound fraction of the tetrahedral antibody was captured by passing the sample through solid-phase polymethyl methacrylate (PMMA) particles (Sapidyne, Cat#2P4510) or 98 micrometer polystyrene particles (Sapidyne, Cat#442178) that had been adsorption-coated with spike protein at a concentration of 30 micrograms/mL.

The captured, unbound fraction was detected using an Alexa Fluor 647-conjugated AffiniPure Goat Anti-Human IgG F(ab’)2 fragment-specific secondary antibody (Jackson ImmunoResearch, Cat#109-605-097) or an AffiniPure Rabbit Anti-Human IgG Fc-gamma fragment-specific secondary antibody (Jackson ImmunoResearch, Cat#309-005-008) at a final concentration of 0.5 micrograms/mL. The KinExA instrument optical path utilized a 620/30 nm bandpass excitation filter, a 670/40 nm bandpass emission filter, and a 645 nm longpass dichroic mirror. Standard equilibrium binding curve analysis was performed using KinExA Pro software v4.3.20 (Sapidyne). Titrant and CBP concentrations utilized in the multi-curve (n-curve) analysis were converted to functional binding-site concentrations based on the calculated valence of the test article, and the final spike trimer concentration was mathematically corrected for biological activity.

## Neutralization of SARS-CoV-2 Spike Protein Binding to ACE2-Expressing Cells

Human embryonic kidney 293T cells stably expressing human ACE2 (293T-hsACE2; Integral Molecular, Cat#C-HA101) were cultured in Dulbecco’s Modified Eagle Medium (DMEM; ATCC, Cat#30-2002) supplemented with 10% fetal bovine serum (FBS; Cytiva, Cat#SH30910.03HI), 10 mM HEPES, 1X penicillin-streptomycin (Thermo Fisher, Cat#15140148), and 1 mg/mL puromycin (Thermo Fisher, Cat#A1113802) in T75 flasks. Prior to the assay, adherent cells were detached using 0.05% Trypsin-EDTA (Thermo Fisher, Cat#25200056), washed with DMEM, and seeded into 96-well V-bottom plates at a density of 5 x 104 cells per well.

Serially diluted tetrahedral spike-binding proteins were added to the cells in duplicate and incubated for 5 min at room temperature. Biotinylated, aggregate-free SARS-CoV-2 spike trimer (ExcellGene) was subsequently added to each well at a final concentration of 2.5 nM, and the mixture was incubated for 1 hour at room temperature. Cells were washed twice with wash buffer composed of PBS containing 1% BSA (Miltenyi Biotec, Cat#130-091-376), resuspended in wash buffer containing Alexa Fluor 488-labeled streptavidin (BioLegend, Cat#405235), and incubated for 1 hour at 4 degrees Celsius. Following two final washes, cells were resuspended in wash buffer and analyzed on a NovoCyte 2000R flow cytometer (Agilent). The mean fluorescence intensity (MFI) of the Alexa Fluor 488 signal corresponding to cell-surface bound biotinylated spike was quantified using NovoExpress Software v1.4.1 (Agilent). Data were analyzed via non-linear regression using GraphPad Prism v5, and the quantity of bound spike trimer was expressed in relative units (RU) based on a standard curve of known spike protein concentrations.

## Live SARS-CoV-2 and NL63 Neutralization Assays

Infectious SARS-CoV-2 (isolate USA-WA1/2020; BEI Resources, Cat#NR-52281) and human coronavirus NL63 (HCoV-NL63; BEI Resources, Cat#NR-470) were propagated via a single passage in Vero/TMPRSS2 cells. Viral stocks were endpoint titrated on Vero/TMPRSS2 cells to determine the median tissue culture infectious dose (TCID₅₀). For neutralization testing, 100 TCID₅₀ of virus per well was pre-incubated for 1 hour at 37 degrees Celsius with serial dilutions of the respective antibody candidate. The virus-antibody mixtures were subsequently inoculated in replicates of eight onto Vero/TMPRSS2 cell monolayers seeded in 96-well plates. Following an incubation period of 7 days at 37 degrees Celsius, the wells were microscopically evaluated and scored for the presence or absence of viral cytopathic effect (CPE).

## Pseudotyped SARS-CoV-2 Neutralization Assays

Neutralization profiles against divergent viral variants were mapped using pseudotyped SARS-CoV-2 reporter virus particles (RVPs) incorporating a *Renilla* luciferase reporter gene (Integral Molecular). Assays evaluated panels representing the following variants and specific mutational footprints: D614G B.1 (Cat#RVP-702L), SARS-CoV-1 Urbani (Cat#RVP-801L), N439K (Cat#RVP-704L), Alpha B.1.1.7 (Cat#RVP-706L), Alpha B.1.1.7 bearing E484K (Cat#RVP-717L), Beta B.1.351 Δ3 (Cat#RVP-724L), Beta B.1.351 (Cat#RVP-707L), Gamma P.1 (Cat#RVP-708L), Delta B.1.617.2 (Cat#RVP-763L), Epsilon B.1.427/B.1.429 (Cat#RVP-713L), Zeta P.2 (Cat#RVP-736L), Eta B.1.525 (Cat#RVP-723L), Iota B.1.526 (Cat#RVP-726L), Kappa B.1.617.1 (Cat#RVP-730L), Lambda C.37 (Cat#RVP-766L), Mu B.1.621 (Cat#RVP-767L), Omicron B.1.1.529 (Cat#RVP-786L), Omicron BA.2 (Cat#RVP-770L), Omicron BA.4/5 (Cat#RVP-744L), Omicron BF.7 (Cat#RVP-779L), Omicron BQ.1.1 (Cat#RVP-782L), Omicron XBB.1.5 (Cat#RVP-786L), and a composite variant combining the N439K substitution with the Beta B.1.351 Δ3 background (Cat#RVP-769L).

Serially diluted anti-spike tetrahedral proteins or monoclonal references were incubated with the pseudotyped virus particles for 1 hour at 37 degrees Celsius. A minimum of nine concentrations were evaluated per protein candidate. Pseudotyped virus incubated in culture medium in the absolute absence of antibody served as the negative control to define 100% infectivity reference values. The equilibrated virus-antibody mixtures were then added to 293T-hsACE2 cells seeded at a density of 2.5 x 10⁵ cells/mL in 96-well plates. Viral entry and reporter expression proceeded for approximately 72 hours at 37 degrees Celsius under a humidified atmosphere containing 5% CO₂.

The resulting bioluminescent signal was quantified using the Renilla-Glo Luciferase Assay System (Promega, Cat#E2710) on a luminometer configured with a 1 ms signal integration time. Relative luminescence units (RLU) derived from the negative control infection wells were normalized and utilized to calculate the percentage neutralization for each experimental antibody concentration. All individual samples were processed in duplicate. Data were analyzed by non-linear regression using GraphPad Prism v9.3.1 to fit a four-parameter logistic curve for the calculation of half-maximal inhibitory concentration (IC₅₀) values.

## Syrian Hamster Efficacy Study

In vivo efficacy evaluation was conducted at BIOQUAL, Inc. (Rockville, MD) utilizing 24 male golden Syrian hamsters (*Mesocricetus auratus*) aged 6 to 8 weeks. Animals were randomized into four experimental groups (n = 6 animals per group). Prophylactic interventions were administered on study day minus one (-1) via intraperitoneal (i.p.) injection. Test articles were formulated in a sterile PBS matrix. The SARS-CoV-2 Beta variant utilized for the infectious challenge (hCoV-19/South Africa/KRISP-K005325/2020; BEI Resources, Cat#NR-54974, Lot#020521-105) carries the following defining amino acid substitutions within the spike glycoprotein relative to the baseline Wuhan-1 reference sequence (NCBI Reference Sequence: NC_045512.2): L18F, D80A, D215A, ΔL242/A243/L244, K417N, E484K, N501Y, D614G, and A701V.

On study day 0, baseline pre-challenge blood samples were collected from all subjects. Animals were subsequently subjected to an intranasal viral challenge comprising 108,750 plaque-forming units (PFU) of the SARS-CoV-2 Beta variant, delivered as a 50 microliter inoculum per nostril. Following the infectious challenge, all animals were clinically monitored twice daily, and individual body weights were recorded daily.

## Pharmacokinetic Assays

In vivo pharmacokinetic parameters were mapped at The Jackson Laboratory (Bar Harbor, ME) utilizing male B6.Cg-Fcgrt Tg(FCGRT)32Dcr/DcrJ mice aged 6 to 8 weeks, which are homozygous for the human neonatal Fc receptor (FcRn) transgene. Baseline body weights were established within 24 hours prior to administration. At time zero (0 hours) on study day 0, each protein therapeutic was administered via a single intravenous (i.v.) bolus injection at a dose of 10 mg/kg adjusted to a dosing volume of 5 mL/kg. Whole-blood samples (25 microliters) were collected via serial bleeding at 5 min, 6 hours, 1 day, 3 days, 5 days, 7 days, 10 days, 14 days, 17 days, 21 days, and 28 days post-injection. Blood samples were immediately transferred into tubes containing 1 microliter of K₃EDTA, processed via centrifugation to isolate plasma, and diluted 1:10 in a matrix of 50% glycerol in PBS. The resulting diluted plasma samples were frozen in 96-well storage plates and maintained at -20 degrees Celsius.

Quantification of circulating therapeutic proteins within the plasma matrices was performed via electrochemiluminescence immunoassay (ECLIA) at Hinge Bio Inc. Recombinant SARS-CoV-2 spike protein (LakePharma, Cat#46328) was diluted to 20 nM in PBS and immobilized onto QuickPlex 96-well plates (Meso Scale Discovery, Cat#L55XA) at a volume of 25 microliters per well. The plates were sealed and incubated overnight at 4 degrees Celsius, followed by blocking with PBS supplemented with 1% BSA (PBS-B) at room temperature for 30 min. Diluted plasma samples (ranging from 250-fold to 4000-fold dilutions in PBS-B) were dispensed into the coated plates using a Biomek i5 liquid handler (Beckman Coulter). The samples were incubated under constant agitation (700 rpm) at room temperature for 60 min.

Target molecules captured on the plates were detected using a biotinylated goat anti-human Fab antibody (Abcam, Cat#ab64666) or a biotinylated goat anti-human ACE2 antibody (R&D Systems, Cat#DY933-05), applied at 25 microliters per well and incubated under constant agitation (700 rpm) at room temperature for 60 min. Plates were washed, and Sulfo-Tag-conjugated streptavidin (Meso Scale Discovery, Cat#R32AD) was introduced as the secondary electrochemiluminescent signaling reagent. Following final plate washes, 150 microliters of MSD Read Buffer A (Meso Scale Discovery, Cat#R92TG) were added immediately prior to plate evaluation on a MESO QuickPlex instrument. Luminescent data were acquired and analyzed using MSD Discovery Workbench v4.0.13 software.

## Pharmacokinetic Analysis

Non-compartmental pharmacokinetic (NCA) parameters were derived from individual plasma concentration-time profiles using Phoenix WinNonlin v8.3 (Certara). Pharmacokinetic constants were calculated for individual mice utilizing the standardized software module designated for intravenous bolus administration. The area under the plasma concentration-time curve (AUC) was calculated via the linear trapezoidal method. The terminal elimination slope (lambda _z) for each subject was derived via linear regression modeling of the terminal data points spanning from approximately day 5 through the final sampling interval at day 28; this terminal elimination constant was subsequently used to calculate the elimination half-life (t_1/2_) and the total AUC extrapolated to infinity (AUC_inf_).

## ACE2 Enzyme Assays

The carboxypeptidase enzymatic activity of the engineered constructs was evaluated by initiating a cleavage reaction containing 0.025 micrograms of wild-type reference protein or 15 micrograms of the engineered ACE2-mutant tetrahedral variant. Substrates evaluated included 0.2 μM angiotensin II (AnaSpec, Cat#AS-20633), bradykinin (AnaSpec, Cat#AS-65642), or apelin-13 (CPC Scientific, Cat#APEL-003). All enzyme kinetics were executed in a specialized reaction buffer composed of PBS supplemented with 10 μM ZnCl₂ maintained at 37 degrees Celsius. Recombinant human ACE2 protein (BioLegend, Cat#79200) was evaluated in parallel as a positive control. Serial reaction aliquots (25 microliters) were collected every 2 min and immediately heat-inactivated at 80 degrees Celsius for 5 min to terminate enzymatic turnover. The absolute concentration of liberated free phenylalanine within the heat-inactivated fractions was quantified using a fluorometric Phenylalanine Assay Kit (Abcam, Cat#ab83376).

## Cell-Surface CD20 and CD19 Binding Assays

Human Toledo B cells (ATCC, Cat#CRL-2631) were cultivated in RPMI-1640 medium (ATCC, Cat#30-2001) supplemented with 10% fetal bovine serum (FBS) and 1 microgram/mL gentamicin (Thermo Fisher Scientific, Cat#15750060) in T25 flasks. For target engagement profiles, cells were distributed into 96-well V-bottom plates at a density of 5 x 10⁴ cells per well in blocking buffer composed of PBS containing 5% normal mouse serum (Millipore Sigma, Cat#M5905). Cells were incubated with 100 nM unlabeled CD20- or CD19-binding therapeutic proteins for 30 min at room temperature.

To evaluate steric occlusion, anti-CD20 (rituximab) and anti-CD19 (FMC63) monoclonal antibodies were chemically conjugated to Alexa Fluor 488 (Abcam, Cat#ab236553) and Alexa Fluor 647 (Abcam, Cat#ab269823), respectively, according to the manufacturer’s protocols. Cells were subsequently incubated with 100 nM of the fluorophore-labeled antibodies for 30 min at room temperature, washed twice with wash buffer composed of PBS containing 0.3% bovine serum albumin (BSA; Millipore Sigma, Cat#A7030), and resuspended in wash buffer containing propidium iodide (Thermo Fisher Scientific, Cat#P3566) to exclude dead cell populations. Flow cytometric acquisition was executed on a NovoCyte 2000R cytometer, and data processing was completed using NovoExpress Software v1.4.1 (Agilent Technologies).

## Antibody-Dependent Cellular Cytotoxicity Assay

Toledo target B cells were maintained in RPMI-1640 medium with GlutaMAX (Thermo Fisher Scientific, Cat#61870036) containing 10% FBS (Cytiva, Cat#SH30070.03IH30-45). To visually differentiate target populations from effector cells, Toledo cells were pre-labeled with 5 micromolar CellTrace Yellow (Thermo Fisher Scientific, Cat#C34567) for 20 min at 37 degrees Celsius. Following metabolic quenching and washing according to the manufacturer’s instructions, 1 x 10³ target cells per well were dispensed in duplicate into 96-well ultra-low attachment microplates (Corning, Cat#7007). Samples were treated with designated CD20- or CD19-binding molecules and 2 micromolar CellEvent Caspase-3/7 Green Detection Reagent (Thermo Fisher Scientific, Cat#C10423) for 10 min at room temperature.

Primary human peripheral blood natural killer (NK) cells (Stemcell Technologies, Cat#70036) were introduced into the wells at a defined effector-to-target (E:T) ratio of 10:1. Fluorogenic caspase-3/7 cleavage kinetics were monitored continuously every hour for 8 hours using an IncuCyte SX5 live-cell imaging system (Sartorius) maintained at 37 degrees Celsius under a humidified atmosphere of 5% CO₂. Apoptotic, caspase-3/7-positive target cells were distinguished from unlabeled apoptotic NK cells via multi-channel fluorescence thresholding. Kinetic datasets were extracted using IncuCyte 2021A software (Sartorius), and the area-under-the-curve (AUC) values (expressed as mean ± s.e.m.) along with non-linear regression profiles were modeled using GraphPad Prism v9.3.1.

## Surface Plasmon Resonance Experiments

High-throughput surface plasmon resonance (SPR) capture kinetic assays were executed on a Carterra LSA single-cycle kinetics instrument (Carterra). Recombinant tetrahedral antibodies or monoclonal references were diluted to 5 micrograms/mL and permanently immobilized onto a Carterra Carboxymethyl Dextran-Hydrogel (CMD-P) sensor chip surface via standard amine coupling chemistry. Recombinant human analytes were prepared in three-fold serial dilution series using a modified HEPES running buffer matrix (10 mM HEPES, 150 mM NaCl, 3 mM EDTA, 0.005% Tween-20, 0.05% n-dodecyl-beta-D-maltoside (DDM), 0.01% cholesteryl hemisuccinate (CHS), pH 7.4, supplemented with 0.05 mg/mL BSA). The concentrated analyte ranges utilized were: FcγRIIIa (2,000 nM to 2.7 nM; R&D Systems, Cat#4325-FC-50), FcγRIIa (R167 variant; 667 nM to 2.7 nM; R&D Systems, Cat#1330-CD-050/CF), FcγRI (55.6 nM to 0.68 nM; R&D Systems, Cat#CF1257-FC), CD19 (500 nM to 0.68 nM; Acrobiosystems, Cat#CD9-H52H2), and complement component C1q (167 nM to 0.68 nM; Complement Technology, Cat#A099).

Recombinant human CD20 protein (167 nM to 0.68 nM; Acrobiosystems, Cat#CD0-H52H3) was diluted independently in an identical three-fold step pattern using the same HEPES running buffer matrix modified with 0.05% DDM and 0.01% CHS to preserve receptor micelle stability. Prepared analytes were injected across the CMD-P sensor surfaces for 5 min during the association phase, followed by an immediate shift to running buffer for an additional 5 min to monitor dissociation kinetics. Complete chip regeneration between successive analyte injection rounds was achieved by passing Pierce IgG Elution Buffer (Thermo Fisher Scientific, Cat#21004) fortified with 1 M NaCl through the flow cells three times for 1 min per interval. Raw binding sensorgrams were processed and fitted to global kinetic models using Carterra Kinetics Software (Carterra).

## Neutralization of Recombinant Human IL-17A and TNF-*α*-Induced IL-6 Secretion

Primary normal human dermal fibroblasts (NHDF; PromoCell, Cat#12302) isolated from adult skin tissue were propagated in fibroblast Growth Medium 2 (PromoCell, Cat#C23020) and seeded overnight into 96-well microplates (PerkinElmer, Cat#6005182) at a density of 2 x 10⁴ cells per well. The following day, the growth media was evacuated and replaced with assay medium containing a pre-established synergistic cytokine formulation consisting of 2.5 ng/mL recombinant human TNF-α (PromoCell, Cat#10291) and 10 ng/mL recombinant human IL-17A (R&D Systems, Cat#7955).

Prior to cell exposure, this synergistic cytokine matrix was pre-incubated for 30 min at 37 degrees Celsius with serially diluted adalimumab, secukinumab, the bispecific anti-TNF-α/anti-IL-17A tetrahedral antibody candidate HB2309, or various defined molar ratios of individual secukinumab and adalimumab controls. Following a 24-hour incubation period, the culture supernatants were harvested, and the absolute concentration of secreted interleukin-6 (IL-6) was quantified via electrochemiluminescence assay using an MSD human IL-6 kit (Meso Scale Discovery, Cat#K151AKB-2) according to the manufacturer’s technical specifications.

## Inhibition of Recombinant Human TNF-*α*-Mediated Cytotoxicity

The neutralizing potencies of the engineered candidates HB2309 and HB2310 were evaluated in an in vitro cytotoxicity assay using the WEHI-164 mouse fibrosarcoma cell line (ATCC, Cat#CRL-1751) expressing endogenous murine TNF-α receptors. Cells were seeded in duplicate at a density of 5 x 10⁴ cells per well into 96-well plates and incubated in standard RPMI-1640 medium supplemented with 10% (v/v) FBS for 18 hours. Serially diluted antibody references or tetrahedral variants (final testing concentration range: 10.2 pM to 60,000 pM) were pre-incubated in medium containing 2 micrograms/mL actinomycin D and 20 IU/mL recombinant human TNF-α for 30 min at 37 degrees Celsius. This mixture was subsequently transferred to the adherent cell monolayers. Following an incubation interval of 20 hours at 37 degrees Celsius, cell viability and metabolic density were analyzed fluorometrically using the CellTiter-Glo Luminescent Cell Viability Assay (Promega, Cat#G7573) according to the manufacturer’s instructions.

## Interleukin-17A Reporter Bioassay

Interleukin-17A signaling cascade neutralization was evaluated using the IL-17 Bioassay Thaw and Use Kit (Promega, Cat#CS2018C08) according to the manufacturer’s instructions. Serial three-fold dilutions of anti-IL-17 compounds, starting from a maximum concentration of 600 nM, were pre-incubated with 20 ng/mL recombinant human IL-17A (PeproTech, Cat#200-17) for 25 min at 37 degrees Celsius. The mixtures were subsequently added to the engineered IL-17 Bioassay Cells, which house a luciferase reporter gene integrated under the control of an endogenous IL-17 pathway-responsive promoter. Residual, non-neutralized IL-17A binds to the surface receptors and drives intracellular downstream activation of luciferase expression. Following a 6-hour incubation period at 37 degrees Celsius, the total luminescent output— proportional to the amount of active, non-neutralized cytokine—was quantified using a SpectraMax Paradigm multi-mode microplate detection platform (Molecular Devices). Raw luminescent data were processed, plotted, and analyzed with GraphPad Prism v9.5.0.

## Non-human primate study

### Virus stock

Vero E6 cells (ATCC CRL-1586) were cultured in Dulbecco’s modified Eagle’s medium (DMEM; Gibco, Darmstadt, Germany) supplemented with 10% fetal calf serum (Gibco). Cells were maintained at 37°C in a humidified atmosphere containing 5% CO_2_. The XBB.1.5 strain was used as described elsewhere (https://pubmed.ncbi.nlm.nih.gov/38214083/). A low-passage (passage 3) virus stock was generated under reduced-serum conditions (2% fetal calf serum), harvested after 4 days, concentrated using Vivaspin 20 mL columns (100 kDa molecular weight cutoff; Sartorius), and titrated by plaque assay on Vero E6 cells. Virus stocks were sequenced to exclude substitutions introduced during isolation and passaging. Cells and virus stocks tested negative for simian virus 5 and mycoplasma contamination.

### Design of non-human primate experiment

All work with infectious SARS-CoV-2 was conducted under BSL-3 conditions at the German Primate Center. Animals were kept in compliance with the regulations of the European Parliament and the Council Directive on the protection of animals used for scientific purposes (2010/63/EU). The experiment was approved by the Lower Saxony State Office for Consumer Protection and Food Safety (approval no.: 33.19-42502-04-20/3601). Eight 4-11 years-old male and female rhesus macaques (*Macaca mulatta*) were randomly assigned to a control group (4 animals) and a prophylaxis group (4 animals). The animals in the prophylaxis group received continuous intravenous (IV) infusion of 100 mg/kg body weight of ACE2-740/B13A one day before infection (day -1). At the day of infection (day 0), all animals were inoculated with SARS-CoV-2 isolate XBB1.5 at 1 x 10^5^ PFU through intranasal and oropharyngeal spray application with small, disposable, mechanic aerosolization devices attached to a 1 mL syringe (MAD nasal device and MADgic drug atomization device for children, Teleflex). For this, 0.25 ml were nebulized into each nostril and 4.5 ml into the trachea. The animals were observed daily for clinical signs. At days 0, 2, and 4 post infection, clinical parameters including bodyweight and body temperature were measured. Blood and nasal and pharyngeal swabs were collected at 3 days before infection (day -3), at day 0 and at days two and four after infection (days 2 and 4). All samples for virological analyses were frozen at -80 °C until further processing. At day 4 post infection, the animals were euthanized using pentobarbital and necropsy was performed.

### TCID_50_ assay

Vero E6 cells were cultured in DMEM with 10% Fetal Calf Serum (FCS) and 1% Penicillin/Streptomycin (PS) in a humidified atmosphere at 37 °C with 5% CO2. Cells were plated at 8 × 10^3^ cells per well in a 96-well plate at 24 h before infection. Cells were then infected with 10-fold serial dilutions of transport media from nose and pharynx swabs samples in 100 µl of DMEM. At 1 hour post-infection (hpi), media was removed, cells were washed with PBS and 100 µl of DMEM was added to the wells. Virus-induced CPE was detected 24 hpi using a Keyence BZ-X8000 microscope. Viral titers (TCID_50_/ml) were calculated using the “Improved Kärber method”. Analysis was performed in quadruplicates.

### Quantitative RT-PCR

Viral RNA was extracted from nose and pharynx swab samples using QIAmp Viral extraction kit (Qiagen) following the manufacturer’s instructions. RT-qPCR was performed adapting an established protocol (PMID: 32235945). Primers and probe targeting SARS-CoV-2 subgenomic E gene were used following the QIAGEN OneStep RT-PCR Kit (Qiagen) protocol. Amplification was performed as follows: reverse transcription 50 °C 30 min, denaturation 95 °C 15 min, followed by 50× cycles of amplification at 95 °C 15 sec, 58 °C 30 sec, 72 °C 20 sec in a thermo cycler (Rotor-gene).

### Quantification of Soluble Analytes Using Multiplex Bead-Based Immunoassay

To simultaneously quantify the concentrations of multiple cytokines and chemokines, a bead-based multiplex sandwich immunoassay was performed using the LEGENDplex™ NHP Inflammation Panel V02 (13-plex) w/ FP (BioLegend, San Diego, CA, USA). Rhesus macaque serum samples were thawed and diluted 1:4 in the provided Assay Buffer. The entire assay procedure, including standard curve preparation, bead incubation, detection of antibody binding, and streptavidin-PE staining, was performed according to the manufacturer’s instructions using a 96-well plate format. Samples were acquired using a Sony ID7000 Spectral Cell Analyzer (Sony Biotechnology, San Jose, CA, USA). Setup beads provided in the kit were used to adjust detector gains and establish proper gating parameters for the target bead populations. Data analysis and quantification were performed using the BioLegend LEGENDplex Data Analysis Software Suite. Analyte concentrations were calculated using a five-parameter logistic (5PL) curve-fitting algorithm. Final concentrations are expressed in pg/mL.

### Statistical analysis

Microsoft Excel (as part of the Microsoft Office software package, version 2019, Microsoft Corporation) and GraphPad Prism 9 version 9.5.1 (GraphPad Software) were used to analyze the data. Statistical significance was set as p values of 0.05 or less. Viral infectivity (TCDI50/ml) and viral load (log10 RNA copies/ml) in swab samples were analyzed by one-way ANOVA (Kruskal-Wallis test) with Dunn’s multiple comparison test.

## Fc binding to Fc*γ*R-expressing CHO cell lines

FcγR-expressing CHO cell lines were obtained from Genscript (Cat#M00586, Cat#M00588 Cat#M00597, Cat#M00598, Cat#M00600, Cat#M00601, Cat#M00602 Genscript, Piscataway, New Jersey) and cultured according to the manufacturer’s recommendations. Before the experiment, cells were trypsinized, counted, and resuspended in the wash buffer (0.3% of BSA, 0.01% NaN3, 1mM EDTA in PBS) in concentration 6.25x10^5^ cells/ml. Forty microliters of cell suspension in the wash buffer were plated in V-bottom plates, resulting in cell concentration 2.5x10^4^ cells/well. Ten microliters of serial diluted test articles were added to respective wells and incubated for 30 minutes on ice. Cells were washed twice and resuspended in wash buffer with 10 ng/ul FITC labeled Goat F(ab’)2 Anti-Human Kappa (Cat#2062-02, Southern Biotech, Birmingham, AL) to human Fab region and incubated for 30 minutes on ice in dark. Cells were washed twice and resuspended in 1% BD Cytofix™ buffer solution (Cat# 554655, BD Bioscience). Incubated 15 minutes, washed and resuspended in wash buffer with LIVE/DEAD NIR (Cat#L10119, Thermofisher Scientific) to exclude dead cell populations. Data were acquired using Agilent Acea Novocyte Flow Cytometer NovoCyte3005 (Cat#2010064) and analyzed with NovoExpress Software v1.4.1 (Agilent Technologies). Mean Fluorescent Intensities for FITC were plotted with GraphPad Prism v9.5.0 software.

## Mathematical Calculation of the Peak Asymmetry Factor (A_s_)

The peak asymmetry factor A_s_ was mathematically evaluated using the standard chromatographic relation: A**_s_** = b/a, where a is the horizontal distance from the peak center perpendicular line to the leading edge of the chromatographic peak, and b is the horizontal distance from the peak center perpendicular line to the trailing edge of the peak. To ensure a mathematically normalized and rigorous comparison completely free of human extraction bias, coordinate extraction was executed utilizing OriginPro’s automated line-tracing algorithm by tracking the exact pixel-intensity centroid, and evaluation boundaries were set to represent an exactly normalized fractional peak height threshold of 6.00% of the respective maximum peak heights (I_max_). Specifically, for the wild-type CLD control shown in **Fig. 6d** (I_max_ = 34,396.55 units), an absolute intensity threshold of 2,064 units was utilized. For the Ala714Cys CLD variant shown in **Fig. 6e** (I_max_ = 38,481.01 units), an absolute intensity threshold of 2,309 units was utilized. For the non-covalent wild-type control evaluated at 6.00% fractional height, the automated coordinates mapped to t_R_ = 13.349 min, x_l_ = 12.846 min, and x_t_ = 14.547 min, yielding width metrics of a = 0.503 min and b = 1.202 min (A**_s_** = 2.38). For the Cys-locked covalent Ala714Cys variant evaluated at 6.00% fractional height, the automated coordinates mapped to t_R_ = 13.159 min, x_l_ = 12.683 min, and x_t_ = 14.121 min, yielding width metrics of a = 0.476 min and b = 0.962 min (A**_s_** = 2.02). This clean contraction of the trailing edge component b under automated line-tracing quantifies the successful structural transition from a continuous dissociation equilibrium to stable zone migration of two discrete, baseline-resolved macromolecular populations.

## Denaturing Intact Mass Spectrometry

Tetrahedral antibodies were evaluated by intact mass spectrometry under non-reducing, denaturing conditions to determine whether they formed non-covalent or Cys-locked CLD dimers. Following removal of N- and O-glycan under non-reducing conditions using Protein Deglycosylation Mix II (NEB, Ipswich, MA), samples were analyzed under denaturing conditions by LS/MS on a Maxis II UHR-QTOF instrument following size-exclusion chromatography (Bruker, Billerica, MA). One to five micrograms of each sample was injected at a flowrate of 20 microliters in a mobile phase consisting of a gradient of 0.1% formic acid in water and 0.1% formic acid in acetonitrile.

## Manuscript preparation

Google Gemini (via Google Workspace Enterprise) was used to review the manuscript text solely for grammar and readability as a final step. No large language model was used for scientific content, data interpretations, or substantive matters.

