## Supplementary Figures and Tables for "Topologically Engineered Tetrahedral Antibody Architectures for Multispecific Therapy"

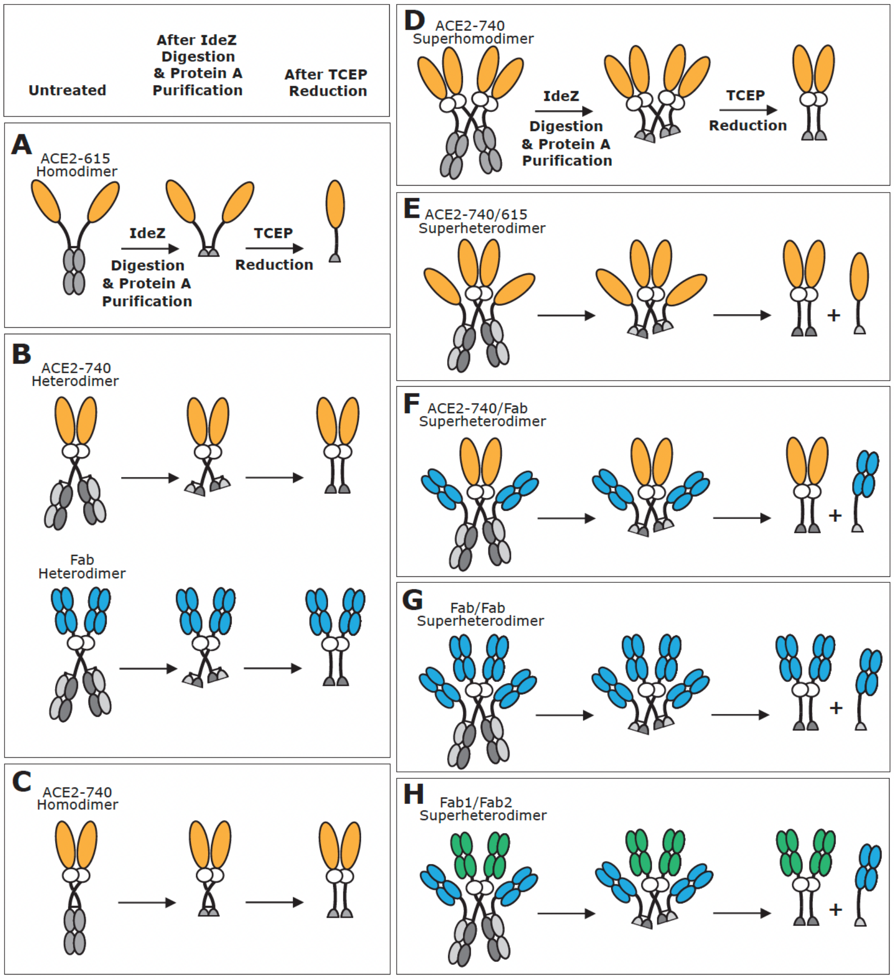

**Supplementary Figure 1**

**Strategy for the confirmation of the structures of the topologically distinct molecules described in these studies using specific cleavage with IdeZ and TCEP reduction.** Molecules were digested with IdeZ protease followed by incubation with protein A beads to remove Fc fragments and incompletely digested molecules. The untreated, IdeZ-treated, and IdeZ-treated/TCEP-reduced molecules were analyzed by SEC/MALS (**Supplementary Figure 2**, **Supplementary Table 1**). Predicted fragments are depicted for **(A)** ACE2-615 dimer (Fig. 1A), **(B)** ACE2-740 tetrahedral antibody (Fig. 1B) and its Fab-substituted counterpart, **(C)** ACE2-740 dimer (Fig. 1C), **(D)** ACE2-740 superdimer (Fig. 1D), **(E)** ACE2-740/615 tetrahedral antibody (Fig. 1E), **(F)** ACE2-740/B13A tetrahedral antibody (Fig. 1F), **(G)** Fab/Fab tetrahedral antibody (Fig. 1G), and **(H)** Fab1/Fab2 tetrahedral antibody (Fig. 1H).

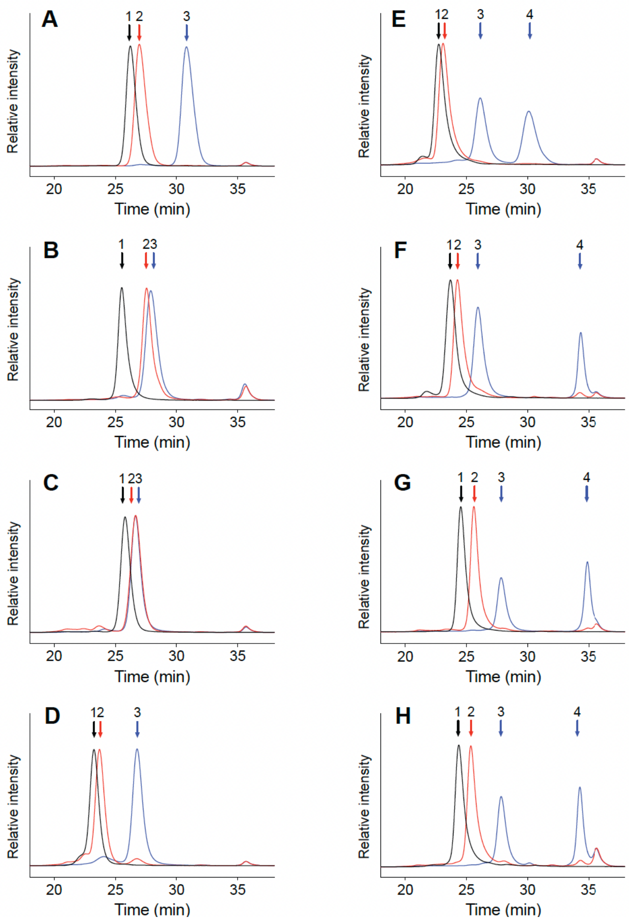

**Supplementary Figure 2**

**SE-HPLC analysis of the topologically distinct molecules described in these studies following IdeZ cleavage and TCEP reduction.** SE-HPLC elution profiles of untreated, IdeZ-treated, and IdeZ-treated/TCEP-reduced molecules are shown by black, red, and blue curves, respectively. Arrow 1 points to the intact molecule, arrow 2 points to the IdeZ cleavage product, and arrows 3 and 4 point to the TCEP-reduced IdeZ cleavage product(s). As expected, IdeZ cleavage products that are dimerized solely by the Fc domain were dissociated upon TCEP reduction of the hinge region interchain disulfides, while IdeZ cleavage products dimerized by the collectrin-like domain did not dissociate upon TCEP reduction. To further confirm each structure, the molar mass of each of the observed peaks was determined by SEC-MALS as described in **Supplementary Table 1**. The following molecules were analyzed: **(A)** ACE2-615 dimer (Fig. 1A), **(B)** Fab tetrahedral antibody (**Supplementary Fig. 1b**), **(C)** ACE2-740 dimer (Fig. 1C), **(D)** ACE2-740 tetrahedral antibody (Fig. 1D), **(E)** ACE2-740/615 tetrahedral antibody (Fig. 1E), **(F)** ACE2-740/B13A tetrahedral antibody (Fig. 1F), **(G)** Fab/Fab tetrahedral antibody (Fig. 1G), and **(H)** Fab1/Fab2 tetrahedral antibody (Fig. 1H).

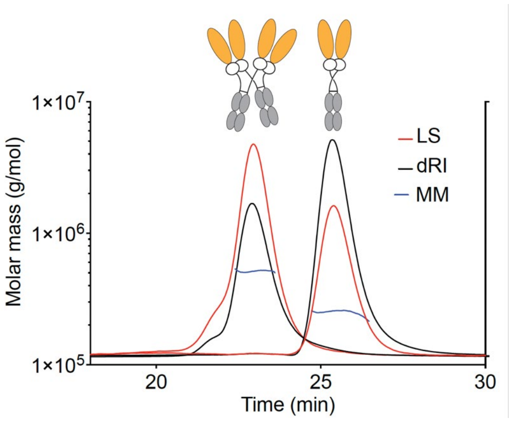

**Supplementary Figure 3**
**SEC-MALS demonstrates that the molar mass of ACE2-740 superdimer is approximately twice that of ACE2-740 dimer.** Purified preparations of ACE2-740 superdimer (Fig. 1C) and ACE2-740 dimer (Fig. 1B) were analyzed individually by SEC-MALS. The profiles are overlaid for comparison. Molar masses are summarized in **Supplementary Table 1**. Abbreviations: LS, light scattering; dRI, differential refractive index; MM, molar mass.

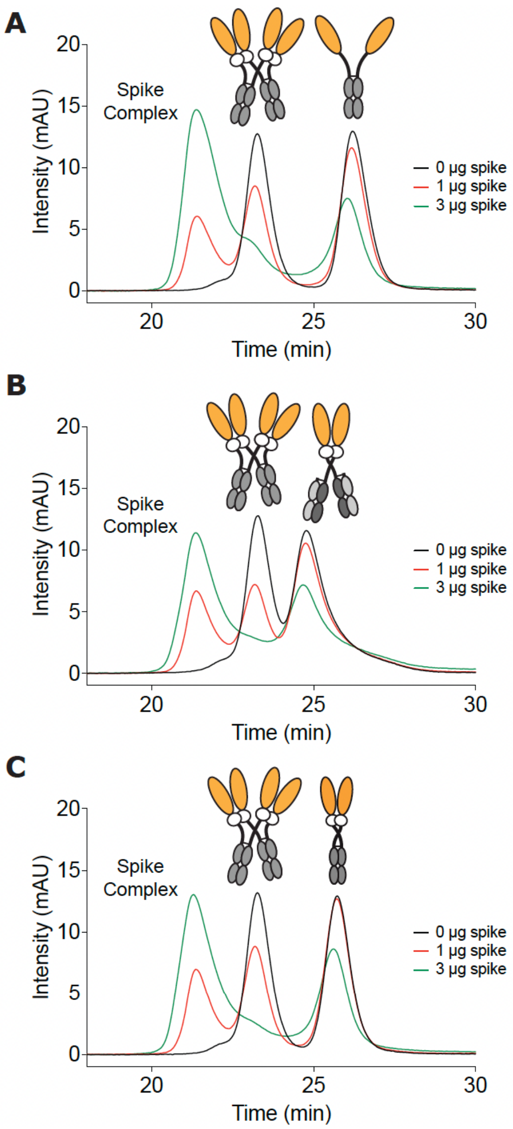

**Supplementary Figure 4**

**Order-of-binding analysis demonstrates that ACE2 superdimer-spike complexes form at the expense of ACE2 dimer-spike complexes.** Titrations with aggregate-free spike trimer were carried out using mixtures of purified ACE2-740 superdimer (Fig. 1C) with purified preparations of the following ACE2 dimers: **(A)** ACE2-615 dimer (Fig. 1A), **(B)** ACE2-740 tetrahedral antibody (Fig. 1D), and **(C)** ACE2-740 dimer (Fig. 1B).

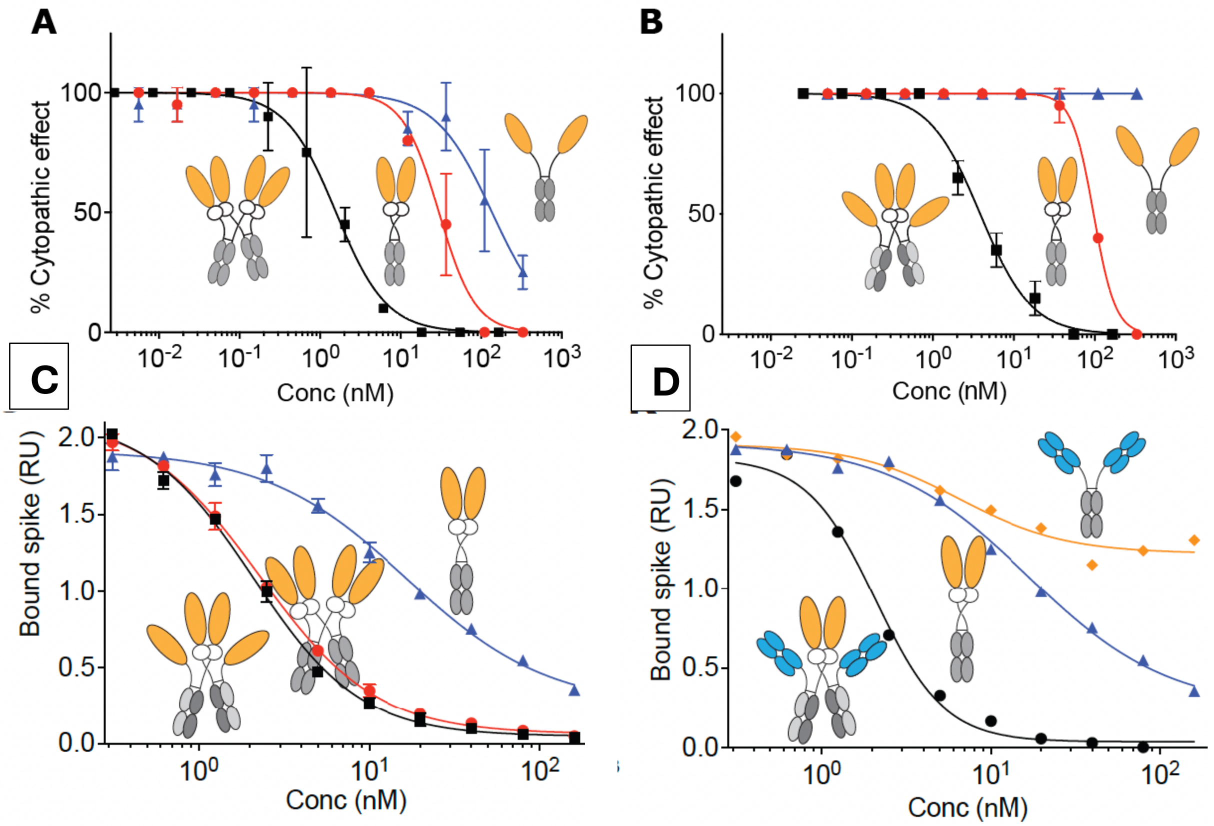

**Supplementary Figure 5**

**Neutralization of live NL63 alphacoronavirus and aggregate free spike trimers**

(**a, b**) Neutralization of live NL63 alphacoronavirus by ACE2 superdimers compared to ACE2 dimers: (**a**) ACE2-740 superdimer (squares), ACE2-740 dimer (circles), ACE2-615 dimer (triangles), and (**b**) ACE2-740/615 tetrahedral antibody (squares), ACE2-740 dimer (circles) ACE2-615 dimer (triangles). (**c, d**) Neutralization of aggregate-free, individual spike trimer binding to cell surface ACE2: (**c**) ACE2-740/615 tetrahedral antibody, ACE2-740 superdimer, and (**d**) ACE2-740 dimer; ACE2-740/B13A superdimer, ACE2-740 dimer, B13A antibody.

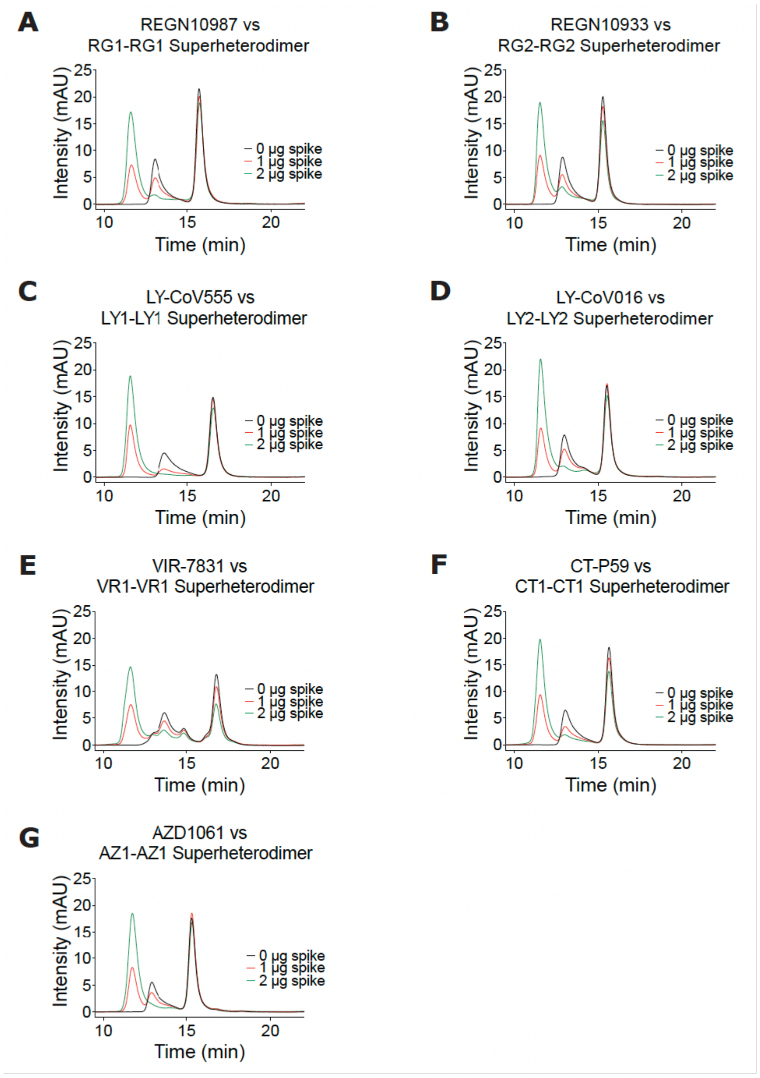

**Supplementary Figure 6**

**Stoichiometric competition binding analysis of antibody Fab/Fab tetrahedral antibodies compared to their parent antibodies.** Titrations with aggregate-free, individual spike trimer were carried out with the following mixtures of antibody Fab/Fab tetrahedral antibodies (Fig. 1G) and their parent antibodies: **(A)** REGN10987 (RG1) and RG1-RG1, **(B)** REGN10933 (RG2) and RG2-RG2, **(C)** LY-CoV555 (LY1) and LY1-LY1, **(D)** LY-CoV016 (LY2) and LY2-LY2, **(E)** VIR-7831 (VR1) and VR1-VR1, **(F)** CT-P59 (CT1) and CT1-CT1, **(G)** AZD1061 (AZ1) and AZ1-AZ1.

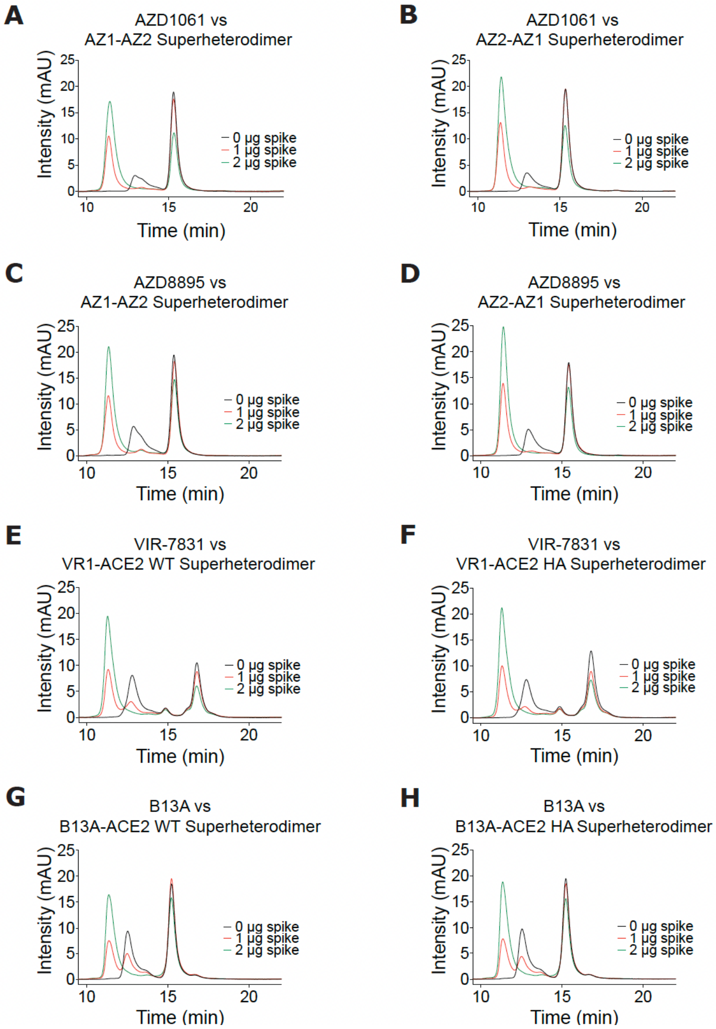

**Supplementary Figure 7**

**Stoichiometric competition binding analysis of bispecific Fab1/Fab2 tetrahedral antibodies and ACE2-740/Fab tetrahedral antibodies compared to their parent antibodies.** Titrations with aggregate-free, individual spike trimer were carried out with the following mixtures of bispecific Fab1/Fab2 tetrahedral antibodies (Fig. 1H) or ACE2-740/Fab tetrahedral antibodies (Fig. 1F) and their parent antibodies: **(A)** AZ1-AZ2 and AZD1061 (AZ1), **(B)** AZ2-AZ1 and AZD1061 (AZ1), **(C)** AZ1-AZ2 and AZD8895 (AZ2), **(D)** AZ2-AZ1 and AZD8895 (AZ2), **(E)** VR1-ACE2 WT and VIR-7831 (VR1), **(F)** VR1-ACE2 HA and VIR-7831 (VR1), **(G)** B13A-ACE2 WT and B13A, and **(H)** B13A-ACE2 WT and B13A. Abbreviations: ACE2 WT, ACE2 with wild-type angiotensin-converting activity; ACE2 HA, ACE2 with H378A mutation which abrogates angiotensin-converting activity.

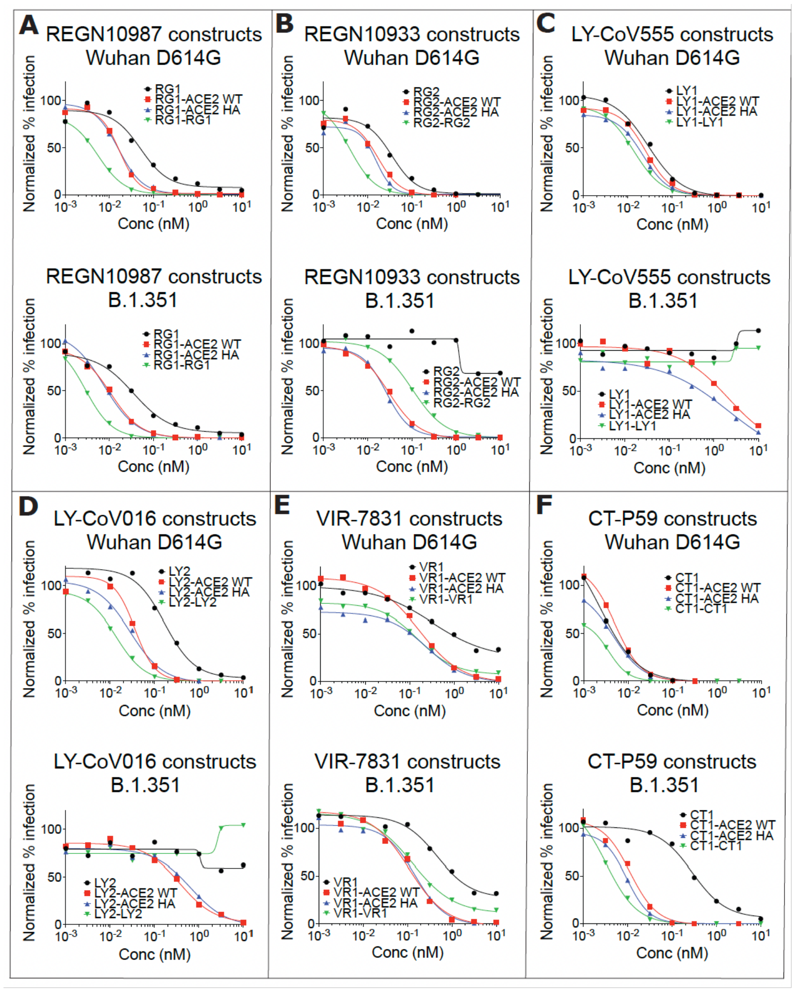

**Supplementary Figure 8**

**Pseudovirus neutralizing activity of ACE2-740/Fab tetrahedral antibodies and antibody Fab/Fab tetrahedral antibodies compared with their parent antibodies.** Neutralization activity against SARS-CoV-2 Wuhan D614G and B.1.351 variants by ACE2-740/Fab tetrahedral antibodies (Fig. 1F), Fab/Fab tetrahedral antibodies (Fig. 1G), and their parent antibodies as follows: **(A)** RG1 (REGN10987), RG1-ACE2 WT, RG1-ACE2 HA, RG1-RG1, **(B)** RG2 (REGN10933), RG2-ACE2 WT, RG2-ACE2 HA, RG2-RG2, **(C)** LY1 (LY-CoV555), LY1-ACE2 WT, LY1-ACE2 HA, LY1-LY1, **(D)** LY2 (LY-CoV016), LY2-ACE2 WT, LY2-ACE2 HA, LY2-LY2, **(E)** VR1 (VIR-7831), VR1-ACE2 WT, VR1-ACE2 HA, VR1-VR1, **(F)** CT1 (CT-P59), CT1-ACE2 WT, CT1-ACE2 HA, CT1-CT1. Abbreviations: ACE2 WT, ACE2 with wild-type angiotensin-converting activity; ACE2 HA, ACE2 with H378A mutation which abrogates angiotensin-converting activity.

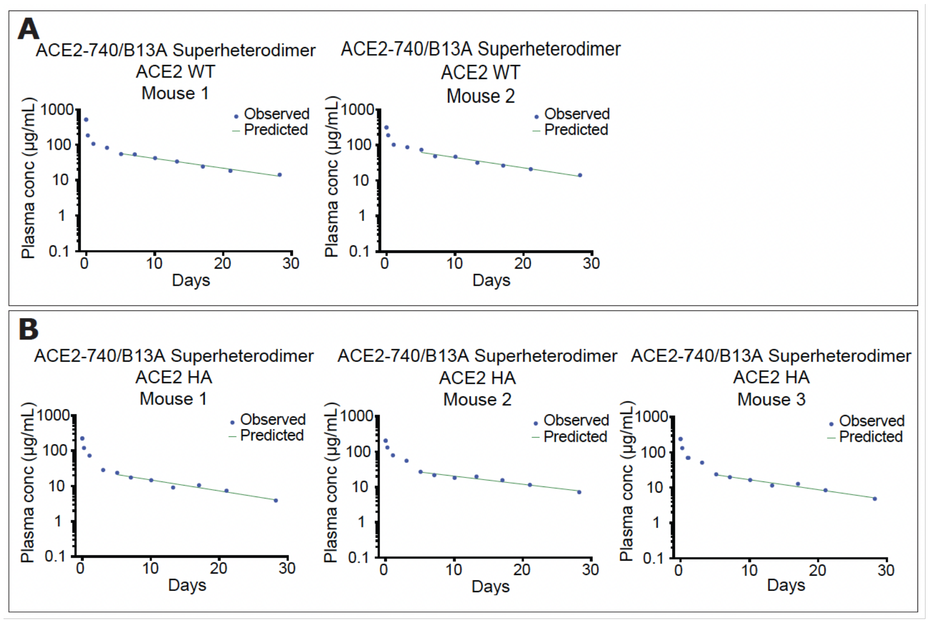

**Supplementary Figure 9**

**Pharmacokinetics of ACE2-740/B13A tetrahedral antibodies in Tg32 mice.** Time course of plasma concentration following a single intravenous administration in Tg32 mice (10 mg/kg) is shown for **(A)** ACE2-740/B13 tetrahedral antibody (ACE2 WT), and **(B)** ACE2-740/B13 HA tetrahedral antibody (ACE2 HA). Plasma concentration was determined by capture with SARS-CoV-2 spike protein and detection with anti-Fab antibody. Comparable results were obtained using anti-ACE2 antibody for detection. Profiles are shown for individual mice. Abbreviations: ACE2 WT, ACE2 with wild-type angiotensin-converting activity; ACE2 HA, ACE2 with H378A mutation which abrogates angiotensin-converting activity.

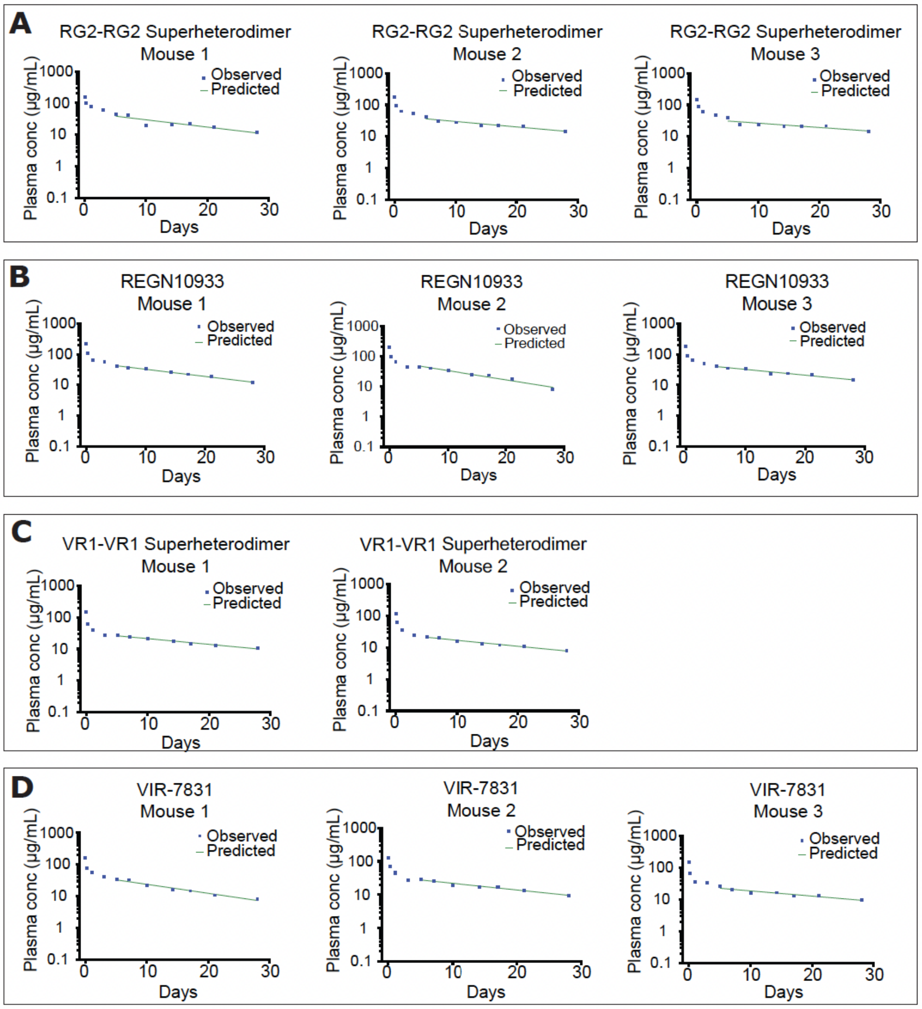

**Supplementary Figure 10**

**Pharmacokinetics of Fab/Fab tetrahedral antibodies in Tg32 mice compared to their parent antibodies.** Time course of plasma concentration following a single intravenous administration in Tg32 mice (10 mg/kg) is shown for **(A)** RG2-RG2 tetrahedral antibody, **(B)** REGN10933 (RG2) antibody, **(C)** VR1-VR1 tetrahedral antibody, **(D)** VIR-7831 (VR1) antibody. Plasma concentration was determined by capturing with SARS-Cov-2 spike protein and detection with anti-Fab antibody. Profiles are shown for individual mice.

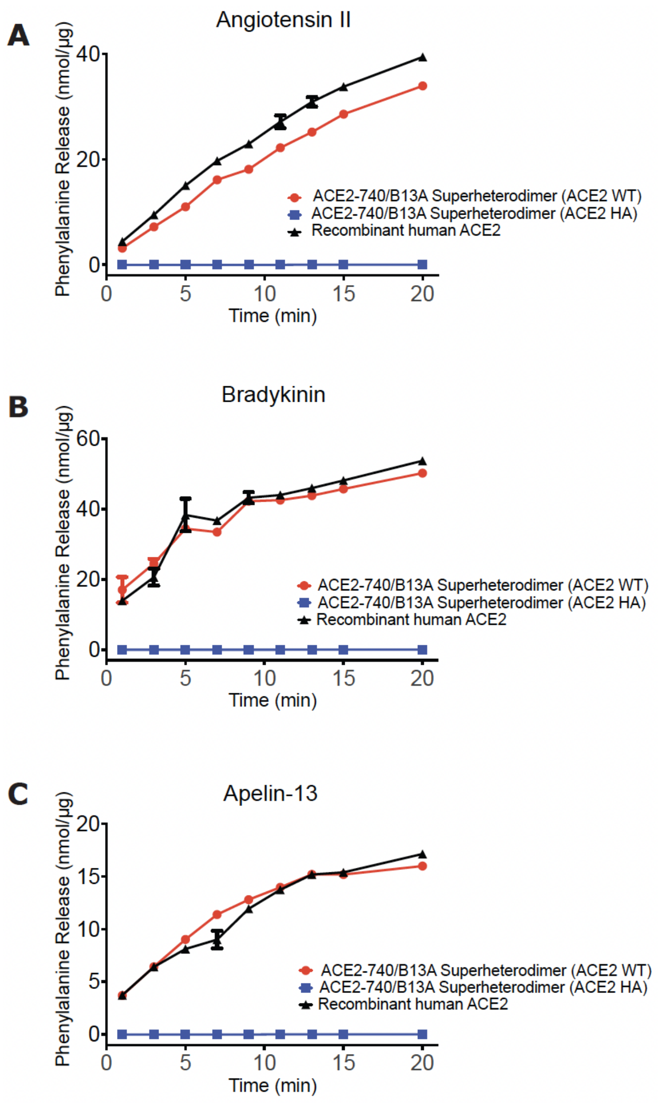

**Supplementary Figure 11**

**Enzymatic activity of ACE2-740/B13A tetrahedral antibodies with wild-type and H378A mutant peptidase domains.** Time course of phenylalanine release by wild-type ACE2-740/B13A tetrahedral antibody (circles), H378A mutant ACE2-740/B13A tetrahedral antibody (squares), and recombinant human ACE2 (triangles) are shown with the following substrates: **(A)** angiotensin II, **(B)** bradykinin, and **(C)** apelin-13.

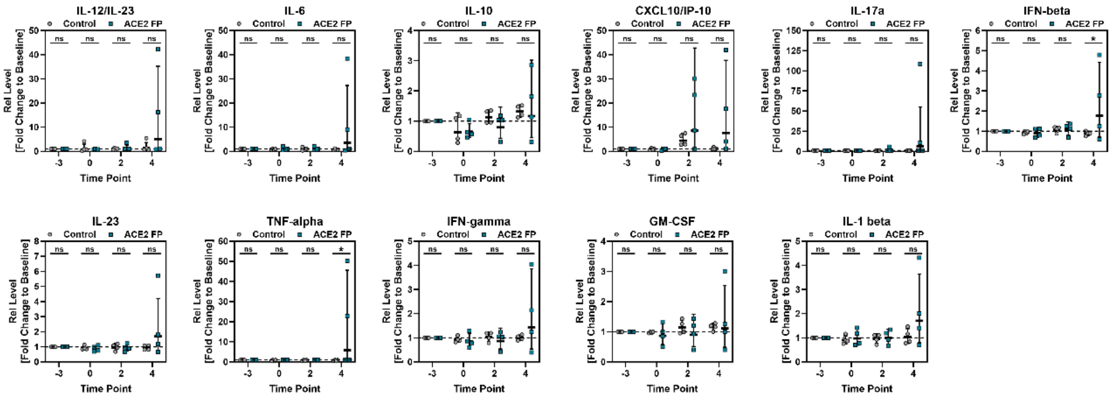

**Supplementary Figure 12**

**Induction of chemokine and cytokine expression upon SARS-CoV-2 infection of rhesus macaques.** Relative changes in the expression of 13 inflammation-associated analytes following SARS-CoV-2 infection. Protein levels were quantified using the LEGENDplex™ NHP Inflammation Panel (13-plex). Data were acquired on a Sony ID7000 spectral analyzer and analyzed with LEGENDplex™ Data Analysis Software using a five-parameter logistic (5PL) curve-fitting algorithm and normalized to baseline (day -3; set as 1). Each symbol represents an individual animal over time with horizontal lines indicating the median. Statistical significance was assessed by two-way analysis of variance (ANOVA) followed by Šídák's multiple-comparisons test.

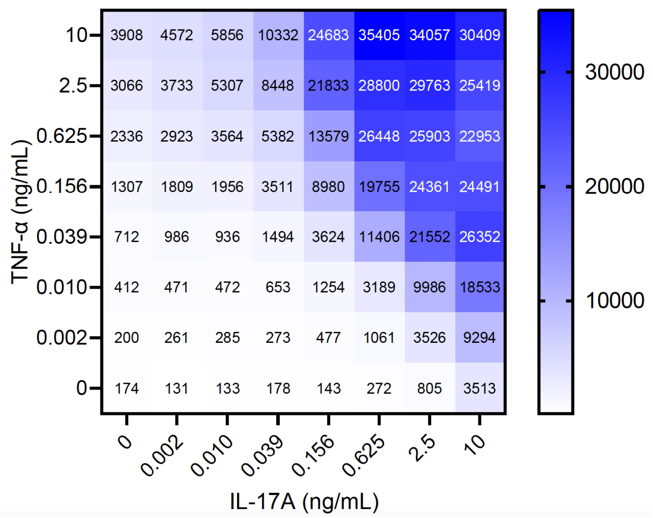

**Supplementary Figure 13**
**Synergistic induction of IL-6 by TNF-α and IL-17A in normal human dermal fibroblasts (NHDF).** Induction of IL-6 is shown for increasing amounts of human TNF-α and IL-17A alone or in combination. IL-6 was measured following a 24-hour cytokine incubation period.

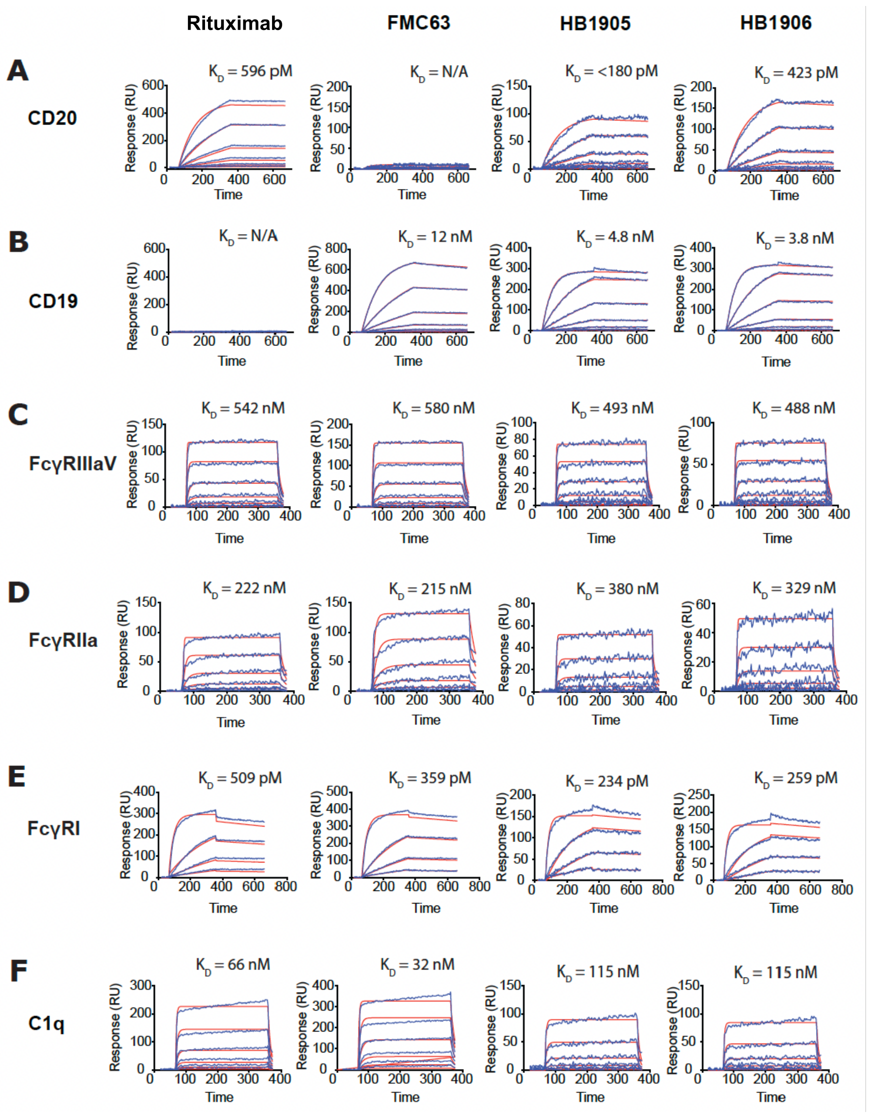

**Supplementary Figure 14**

**Surface plasmon resonance (SPR) binding analysis of anti-CD20/anti-CD19 bispecific Fab1/Fab2 tetrahedral antibodies to CD20, CD19, Fc gamma receptors, and complement C1q.** SPR binding by HB1905 and HB1906 anti-CD20/CD19 bispecific tetrahedral antibodies (Fig. 1H) and rituximab and FMC63 parent antibodies is shown for the following monovalent molecular targets: **(A)** CD20, **(B)** CD19, **(C)**, Fc gamma RIIIa (CD16a) (Val158), **(D)** Fc gamma RIIa (CD32) (Arg131), **(E)** Fc gamma RI (CD64), and **(F)** complement component C1q.

| Structure | Untreated | | IdeZ-treated | | IdeZ-treated/TCEP-reduced | | | |
|---|---|---|---|---|---|---|---|---|
| | Peak 1 Molar Mass | | Peak 2 Molar Mass | | Peak 3 Molar Mass | | Peak 4 Molar Mass | |
| | Experimental (kDa) | Theoretical (kDa) | Experimental (kDa) | Theoretical (kDa) | Experimental (kDa) | Theoretical (kDa) | Experimental (kDa) | Theoretical (kDa) |
| Fig. 1A | 213.3 | 212.5 | 163.9 | 162.0 | 82.7 | 81.0 | -- | -- |
| Fig. S1B | 245.3 | 240.4 | 144.7 | 139.6 | 139.6 | 137.4 | -- | -- |
| Fig. 1C | 248.3 | 246.7 | 194.7 | 196.3 | 195.7 | 196.3 | -- | -- |
| Fig. 1D | 493.1 | 493.4 | 384.8 | 392.6 | 191.6 | 196.3 | -- | -- |
| Fig. 1E | 458.8 | 460.2 | 356.2 | 359.4 | 196.6 | 196.5 | 78.8 | 81.5 |
| Fig. 1F | 395.3 | 392.2 | 296.1 | 291.4 | 203.3 | 196.5 | 47.3 | 47.5 |
| Fig. 1G | 321.3 | 331.6 | 225.3 | 230.8 | 127.5 | 135.4 | 47.2 | 47.7 |
| Fig. 1H | 328.6 | 331.6 | 234.9 | 230.8 | 138.9 | 135.6 | 48.9 | 47.6 |

**Supplementary Table 1. Molar masses determined for the topologically distinct molecules described in these studies and their IdeZ cleavage products confirms their structures.** The molar mass of each of the observed peaks in **Supplementary Figure 2** was determined by SEC-MALS. The theoretical molar mass of each of the observed peaks in **Supplementary Figure 2** was calculated using the known amino acid sequence of each of the predicted structures depicted in **Supplementary Figure 1**, together with the known N-glycan and O-glycan modifications of each of the predicted structures.[30]

| Structure | Molecule | SARS-CoV-2 | | |
|---|---|---|---|---|
| | | IC50 (nM) | Relative Potency | Relative Potency |
| Fig. 1C | ACE2-740 homodimer | 17.2 | 1 | |
| Fig. 1D | ACE2-740 superhomodimer | 1.93 | 8.9 | |
| Fig. 1E | ACE2-740/615 superheterodimer | 1.62 | 10.6 | |
| Fig. 1C | ACE2-740 homodimer | 17.2 | | 1 |
| mAb | B13 antibody | 5.91 | | 2.9 |
| Fig. 1F | ACE2-740/B13A superheterodimer | 1.54 | | 11.1 |

**Supplementary Table 2. Neutralization of SARS-CoV-2 spike trimer binding to cell surface ACE2 receptors.** Neutralization activity ($IC_{50}$) against binding of aggregate-free spike trimer to ACE2 receptors expressed on the surface of 293 embryonic kidney cells is shown for the following molecules: (Fig. 2G) ACE-740 dimer, ACE2-740 superdimer, ACE2-740/615 tetrahedral antibody; (Fig. 2H) ACE-740 dimer, B13A antibody, ACE2-740/B13A tetrahedral antibody.

| Structure | Molecule | SARS-CoV-2 Variant | Fab Type | | | | | |
|---|---|---|---|---|---|---|---|---|
| | | | REGN10987 IC50 (nM) | REGN10933 IC50 (nM) | LY-CoV555 IC50 (nM) | LY-CoV016 IC50 (nM) | VIR-7831 IC50 (nM) | CT-P59 IC50 (nM) |
| mAb | Parent antibody | D614G | 0.051 | 0.035 | 0.029 | 0.168 | 0.318 | 0.002 |
| Fig. 1F | ACE2-740/Fab superheterodimer (ACE2 WT) | D614G | 0.017 | 0.017 | 0.026 | 0.036 | 0.165 | 0.005 |
| Fig. 1F | ACE2-740/Fab superheterodimer (ACE2 HA) | D614G | 0.016 | 0.016 | 0.022 | 0.029 | 0.249 | 0.005 |
| Fig. 1G | Fab/Fab superheterodimer | D614G | 0.005 | 0.004 | 0.013 | 0.013 | 0.153 | 0.004 |
| mAb | Parent antibody | B.1.135 | 0.035 | >10 | >10 | >10 | 0.466 | 0.286 |
| Fig. 1F | ACE2-740/Fab superheterodimer (ACE2 WT) | B.1.135 | 0.011 | 0.030 | 2.104 | 0.371 | 0.108 | 0.011 |
| Fig. 1F | ACE2-740/Fab superheterodimer (ACE2 HA) | B.1.135 | 0.008 | 0.026 | 2.131 | 0.651 | 0.148 | 0.010 |
| Fig. 1G | Fab/Fab superheterodimer | B.1.135 | 0.003 | 0.103 | >10 | >10 | 0.129 | 0.003 |

**Supplementary Table 3. Pseudovirus neutralizing activity of ACE2-740/Fab tetrahedral antibodies and antibody Fab/Fab tetrahedral antibodies compared with their parent antibodies.** Neutralization activity ($IC_{50}$) against Wuhan D614G and B.1.351 variants by ACE2-740/Fab tetrahedral antibodies, Fab/Fab tetrahedral antibodies, and their parent antibodies. Parent antibodies: RG1 (REGN10987), RG2 (REGN10933), LY1 (LY-CoV555), LY2 (LY-CoV016), VR1 (VIR-7831), and CT1 (CT-P59); ACE2-740/Fab tetrahedral antibodies (ACE2 WT): RG1-ACE2 WT, RG2-ACE2 WT, LY1-ACE2 WT, LY2-ACE2 WT, VR1-ACE2 WT, CT1-ACE2 WT; ACE2-740/Fab tetrahedral antibodies (ACE2 HA): RG1-ACE2 HA, RG2-ACE2 HA, LY1-ACE2 HA, LY2-ACE2 HA, VR1-ACE2 HA, CT1-ACE2 HA; Fab/Fab tetrahedral antibodies: RG1-RG1, RG2-RG2, LY1-LY1, LY2-LY2, VR1-VR1, CT1-CT1. Abbreviations: ACE2 WT, ACE2 with wild-type angiotensin-converting activity; ACE2 HA, ACE2 with the H378A mutation which abrogates angiotensin-converting activity.

| | Molecule | Pseudovirus Neutralizing Activity IC50 (nM) | | | | | | | | | | | | | | | | | | | | |
|---|---|---|---|---|---|---|---|---|---|---|---|---|---|---|---|---|---|---|---|---|---|---|
| | | Pre-Omicron Strains | | | | | | | | | | | | | | | Omicron Strains | | | | | |
| | | SARS-CoV-1 Urbani | D614G B.1 | N439K | Alpha B.1.1.7 | Alpha B.1.1.7 E484K | Beta B.1.351 | Gamma P.1 | Delta B.1.617.2 | Epsilon B.1.427/ B.1.429 | Zeta P.2 | Eta B.1.525 | Iota B.1.526 | Kappa B.1.617.1 | Lambda C.37 | Mu B.1.621 | Omicron B.1.1.529 | Omicron BA.2 | Omicron BA.4/5 | Omicron BF.7 | Omicron BQ.1.1 | Omicron XBB.1.5 |
| | **HB1507** | 1.138 | 0.242 | 0.025 | 0.081 | 0.049 | 0.059 | 0.041 | 0.100 | 0.118 | 0.101 | 0.085 | 0.162 | 0.132 | 0.102 | 0.080 | 0.016 | 0.037 | 0.036 | 0.067 | 0.488 | 0.101 |
| | **HB1516** | 0.123 | 0.048 | 0.126 | 0.032 | 0.013 | 0.023 | 0.031 | 0.015 | 0.013 | 0.028 | 0.030 | 0.027 | 0.017 | 0.043 | 0.016 | 0.990 | 0.998 | 0.794 | 2.029 | 3.205 | 2.543 |
| Authorized Antibodies | **REGN10987** | >10 | 0.030 | 1.285 | 0.016 | 0.012 | 0.032 | 0.012 | 0.059 | 0.033 | 0.033 | 0.045 | 0.075 | 0.048 | 0.190 | 0.025 | >10 | >10 | ND | ND | ND | ND |
| | **REGN10933** | >10 | 0.045 | 0.027 | 0.023 | 0.746 | 2.374 | 2.538 | 0.011 | 0.022 | 0.469 | 0.446 | 0.895 | 0.385 | 0.020 | 0.637 | >10 | >10 | ND | ND | ND | ND |
| | **LY-CoV555** | >10 | 0.042 | 0.039 | 0.034 | >10 | >10 | >10 | >10 | >10 | >10 | >10 | >10 | >10 | >10 | >10 | >10 | >10 | ND | ND | ND | ND |
| | **LY-CoV016** | >10 | 0.271 | 0.176 | 2.471 | >10 | >10 | >10 | 0.055 | 0.217 | 0.352 | 0.336 | 0.875 | 0.162 | 0.054 | 3.148 | >10 | >10 | ND | ND | ND | ND |
| | **AZD1061** | >10 | 0.050 | 0.060 | 0.036 | 0.039 | 0.046 | 0.033 | 0.119 | 0.082 | 0.045 | 0.029 | 0.050 | 0.166 | 0.162 | 0.164 | >10 | 0.041 | ND | ND | ND | ND |
| | **AZD8895** | >10 | 0.033 | 0.019 | 0.042 | 0.786 | 0.240 | 0.078 | 0.020 | 0.021 | 0.212 | 0.282 | 0.384 | 0.039 | 0.007 | 0.229 | >10 | >10 | ND | ND | ND | ND |
| | **VIR-7831** | 0.468 | 0.461 | 0.396 | 0.619 | 0.281 | 0.354 | 0.371 | 0.552 | 0.351 | 0.471 | 0.635 | 0.232 | 0.440 | 0.943 | 0.514 | 2.171 | >10 | ND | ND | ND | ND |
| | **CT-P59** | >10 | 0.007 | 0.005 | 0.019 | 0.574 | 1.085 | 0.593 | 0.258 | 0.223 | 0.029 | 0.015 | 0.044 | 0.137 | 0.106 | 0.284 | >10 | >10 | ND | ND | ND | ND |
| | **B13A** | 0.110 | 0.068 | 0.056 | 0.076 | 0.035 | 0.202 | 0.070 | 0.031 | 0.012 | 0.129 | 0.216 | 0.108 | 0.033 | 0.272 | 0.011 | >10 | >10 | ND | ND | ND | ND |

**Supplementary Table 4. Pseudovirus neutralizing activity of ACE2-740/615 (HB1507) and ACE2-740/B13A (HB1516) tetrahedral antibodies compared with eight clinically authorized antibodies.** Neutralization activity ($IC_{50}$) of HB1507 (Fig. 1E), and HB1516 (Fig. 1F), compared with eight clinically authorized antibodies (REGN10987, REGN10933, LY-CoV555, LY-CoV016, AZD1061, AZD8895, VIR-7831, CT-P59) and the preclinical antibody B13A. Results for seventeen major SARS-CoV-2 variants are shown: Alpha B.1.1.7, Beta B.1.351, Gamma P.1, Delta B.1.617.2, Epsilon B.1.427/B.1.429, Zeta P.2, Eta B.1.525, Iota B.1.526, Kappa B.1.617.1, Lambda C.37, Mu B.1.621, Omicron B.1.1.529 Omicron BA.2, Omicron BA.4/5, Omicron BF.7, Omicron BQ.1.1 and Omicron XBB.1.5. Results are also shown for SARS-CoV-2 variants Wuhan D614G, N439K, and Alpha B.1.1.7 E484K, and the SARS-CoV-1 Urbani variant. The HB1507 and HB1516 tetrahedral antibodies have the ACE2 H378A mutation which abrogates angiotensin-converting activity and the P329G, L234A, L235A triple mutation which abrogates Fc receptor binding activity.

| Structure | Protein ID | Molecule | N439K | | B.1.351 | | N439K/B.1.351 | |
|---|---|---|---|---|---|---|---|---|
| | | | IC50 (nM) | Relative Potency | IC50 (nM) | Relative Potency | IC50 (nM) | Relative Potency |
| Fig. 1H | HB1701 | RG1-RG2 superheterodimer | 0.011 | 1.5 | 0.008 | 2.7 | 0.051 | 25.3 |
| | HB1705 | RG2-RG1 superheterodimer | 0.021 | 0.8 | 0.030 | 0.7 | 0.074 | 17.5 |
| | HB1713 | RG1-RG2 superheterodimer | N.D. | N.D. | N.D. | N.D. | 0.057 | 22.6 |
| | HB1722 | RG2-RG1 superheterodimer | 0.008 | 2.0 | 0.009 | 2.2 | 0.080 | 16.2 |
| mAb | REGN10987 | REGN10987 | 4.437 | 0.0 | 0.021 | 1.0 | 3.111 | 0.4 |
| | REGN10933 | REGN10933 | 0.019 | 0.9 | 6.740 | 0.0 | 9.322 | 0.1 |
| | REGN-COV2 | REGN10987+REGN10933 | 0.016 | 1.0 | 0.020 | 1.0 | 1.297 | 1.0 |
| Fig. 1E | HB1507 | ACE2-740/615 superheterodimer | 0.108 | 0.2 | 0.059 | 0.3 | 0.034 | 38.5 |
| | HB1553 | ACE2-740/615 superheterodimer | 0.126 | 0.1 | 0.090 | 0.2 | 0.023 | 57.0 |
| Fig. 1F | HB1515 | ACE2-740/B13A superheterodimer | 0.029 | 0.6 | 0.027 | 0.8 | 0.029 | 45.2 |
| | HB1516 | ACE2-740/B13A superheterodimer | 0.025 | 0.6 | 0.023 | 0.9 | 0.025 | 52.8 |

**Supplementary Table 5. Pseudovirus neutralizing activity of tetravalent bispecific (Fab1/Fab2) tetrahedral antibodies compared with two-antibody cocktails of their parent antibodies.** Pseudovirus neutralization activity ($IC_{50}$) against the N439K, B.1.351, and N439K/B.1.351 variants is shown for: tetravalent bispecific RG1-RG2 tetrahedral antibodies (HB1701, HB1722) and RG2-RG1 tetrahedral antibodies (HB1705, HB1722), compared against the REGN10987 (RG1) and REGN10933 (RG2) parent antibodies used as single agents or as a two-antibody cocktail (REGN-COV2). Neutralization activity is also shown for tetravalent monospecific ACE2-740/615 tetrahedral antibodies (HB1507, HB1553), and tetravalent bispecific ACE2-740/B13A tetrahedral antibodies (HB1515, HB1516).

| Molecule | Specific Activity (pmol/min/μg) | | |
|---|---|---|---|
| | Angiotensin II | Bradykinin | Apelin-13 |
| ACE2-740/B13A superheterodimer (ACE2 WT) | 1,655.20 | 1,724.80 | 688.00 |
| ACE2-740/B13A superheterodimer (ACE2 HA) | 0.76 | 1.14 | 1.01 |
| Recombinant human ACE2 | 1,874.40 | 2,032.40 | 764.40 |

**Supplementary Table 6. Specific activity of ACE2-740/B13A tetrahedral antibodies with wild-type and H378A mutant peptidase domains.** Specific activity of ACE2-740/B13A tetrahedral antibody (ACE2 WT), ACE2-740/B13A tetrahedral antibody (ACE2 HA), and recombinant human ACE2 as measured by phenylalanine release with the following substrates: **(A)** angiotensin II, **(B)** bradykinin, and **(C)** apelin-13. Abbreviations: ACE2 WT, ACE2 with wild-type angiotensin-converting activity; ACE2 HA, ACE2 with the H378A mutation which abrogates angiotensin-converting activity.

| Structure | Configuration | Protein ID | Specificity | WEHI-164 TNF-α Cytotoxicity Assay | | IL-17A Reporter Assay | |
|---|---|---|---|---|---|---|---|
| | | | | IC50 (nM) | Relative Potency | IC50 (nM) | Relative Potency |
| mAb | Ad | Adalimumab | anti-TNF-α | 0.061 | 1.0 | > 600 | N/A |
| mAb | Se | Secukinumab | anti-IL-17A | > 60 | N/A | 15.03 | 1.0 |
| Fig. 1H | Ad x Se | HB2309 | Bispecific anti-TNF-α/anti-IL-17A | 0.087 | 0.7 | 1.26 | 12.0 |
| Fig. 1H | Ad x Se | HB2310 | Bispecific anti-TNF-α/anti-IL-17A | 0.091 | 0.7 | 1.66 | 9.1 |
| Fig. 1H | Se x Ad | HB2313 | Bispecific anti-TNF-α/anti-IL-17A | 0.059 | 1.0 | 13.81 | 1.1 |
| Fig. 1H | Se x Ad | HB2314 | Bispecific anti-TNF-α/anti-IL-17A | 0.059 | 1.0 | 14.45 | 1.0 |

**Supplementary Table 7. Neutralizing activity of anti-TNF-α/anti-IL-17A bispecific Fab1/Fab2 tetrahedral antibodies compared against their parent antibodies adalimumab and secukinumab.** Neutralizing activity against TNF-α and IL-17A shown for bispecific tetrahedral antibodies HB2309 and HB2310 (Ad x Se configuration), and HB2313 and HB2314 Se x Ad configuration) compared against their parent antibodies. Neutralization activity (IC50) was determined for TNF-α-induced cytotoxicity in WEHI-164 cells, and for IL-17A activity in an IL-17A reporter assay.

| Structure | Protein ID | Specificity | Plate 1 | | Plate 2 | |
|---|---|---|---|---|---|---|
| | | | EC50 (pM) | Relative Potency | EC50 (pM) | Relative Potency |
| mAb | Rituximab | anti-CD20 | 177 | 1.0 | 135.5 | 1.0 |
| mAb | FMC63 | anti-CD19 | 5.0 | 35.4 | 5.0 | 27.1 |
| mAb + mAb | Rituximab + FMC63 | anti-CD20 + anti-CD19 | ND | ND | 78.8 | 1.7 |
| Fig. 1H | HB1905 | Bispecific anti-CD20/anti-CD19 | 2.0 | 88.4 | ND | ND |
| Fig. 1H | HB1906 | Bispecific anti-CD20/anti-CD19 | 1.8 | 98.2 | ND | ND |

**Supplementary Table 8. ADCC activity of anti-CD20/anti-CD19 bispecific Fab1/Fab2 tetrahedral antibodies HB1905 and HB1906 compared with their parent antibodies.** ADCC activity ($EC_{50}$) against CD20/CD19-positive Toledo cells is shown for HB1905 and HB1906 and their parent antibodies rituximab (anti-CD20) and FMC63 (anti-CD19) used as single agents or as a two-antibody cocktail.
